# Diversity in recombination hotspot characteristics and gene structure shape fine-scale recombination patterns in plant genomes

**DOI:** 10.1101/2023.12.12.571209

**Authors:** Thomas Brazier, Sylvain Glémin

## Abstract

During the meiosis of many eukaryote species, crossovers tend to occur within narrow regions called recombination hotspots. In plants, it is generally thought that gene regulatory sequences, especially promoters and 5’-3’ untranslated regions, are enriched in hotspots, but this has been characterized in a handful of species only. We also lack a clear description of fine-scale variation in recombination rates within genic regions and little is known about hotspot position and intensity in plants. To address this question we constructed fine-scale recombination maps from genetic polymorphism data and inferred recombination hotspots in eleven plant species. We detected gradients of recombination both in 5’ and 3’ of genic regions in most species, yet gradients varied in intensity and shape depending on specific hotspot locations and gene structure. To further characterize recombination gradients, we decomposed them according to gene structure by rank and number of exons. We generalized the previously observed pattern that recombination hotspots are organized around the boundaries of coding sequences, especially 5’ promoters. However, our results also provided new insight into the relative importance of the 3’ end of genes in some species and the possible location of hotspots away from genic regions in some species. Variation among species seemed driven more by hotspot location among and within genes than by differences in size or intensity among species. Our results shed light on the variation in recombination rates at a very fine scale, more detailed than whole genome averaged estimates used so far, revealing the diversity and complexity of genic recombination gradients emerging from the interaction between hotspot location and gene structure.

## Introduction

Meiotic recombination is a general feature of sexually reproducing species. During meiosis, crossovers (COs) ensure proper segregation of homologous chromosomes and reshuffle alleles from the two parental genomes. The distribution of COs is not homogeneous along chromosomes and varies at large and short scales [1–4]. Characterizing and understanding where recombination occurs have many implications because recombination has both local molecular effects (e.g. mutagenic effects, gene conversion) and indirect effects through the breakdown of linkage disequilibrium [5].

In many species described so far, recombination tends to occur within narrow regions called recombination hotspots [6–9], but in some species, such as *Drosophila* and *Caenorhabditis* species, proper hotspots are lacking. When they exist, the general view is that two main mechanisms determine where hotspots occur across the genome. In many mammals, the location of recombination hotspots is directed by the PRDM9 protein [10, 11]. The zinc-finger domain that binds to specific DNA motifs is evolving fast, thus changing recombination landscapes over short evolutionary times [12, 13]. The PRDM9 gene is believed to be conserved across several vertebrate lineages because it redirects recombination away from the genic region to avoid deleterious mutations and chromosomal rearrangements [10, 14, 15]. Recently, PRDM9-driven hotspots have also been identified in non-mammalian species [16]. In contrast, many animals but also plants and fungi, do not have a PRDM9-like system [14]. In such species where PRDM9 is lacking, recombination was found to occur preferentially in gene regulatory sequences, especially in promoters and 5’-3’ untranslated regions (UTRs) [17–22]. Because this was found in very phylogenetically divergent species such as plants, yeasts and birds [19, 23, 24] and when PRDM9 is experimentally knocked-out [18], it is thought to be the ancestral mechanism for locating recombination hotspots. Mechanistically, recombination hotspots seem globally associated with nucleosome occupancy and methylation patterns that define flanking regions of genes [4, 19, 25]. Although recombination hotspots can overlap coding sequences, both mechanisms tend to lower recombination rates within genes compared to flanking regions and/or non-genic regions.

When hotspots are targeted to promoter like features it can generate recombination gradients within genes, as observed in a few plant species [26–30]. In rice, recombination hotspots overlap genes between the Transcription Starting Site (TSS) and the Transcription Termination Site (TTS), potentially increasing recombination in the first and last exons [30]. In *Mimulus guttatus*, recombination is higher in the first exon than in 5’ noncoding sequences [29]. Even if recombination hotspots strictly remain in flanking regions, recombination tracts span hundreds of base pairs and are likely to end within genes. Indeed, 5’ ends of genes experience gene conversion gradients [31–33] and the combination of GC-biased gene conversion (gBGC) and 5’-3’ recombination gradients would provide an explanation for GC gradients along genes commonly observed among plant species [34–37]. Therefore actual GC gradients are indirect evidence that recombination gradients could be widespread and organized as a function of the number of exons in addition to the distance to TSS and TTS, as observed for GC content [35, 36].

However, this simple dichotomous view has been recently challenged by the exploration of recombination landscapes in various non model species. In mammals, human and mouse, where PRDM9 mechanism was initially identified and characterized in detail, appear as extremes in a more continuous distribution where PRDM9 hotspot could co-occur with PRDM9-independent hotspots directed to promoter regions [38]. Such co-occurrence of PRDM9 and promoter hotspots has also be observed in a snake species [39]. Similarly, we do not know whether the precise location of hotspots around promoter features is the same and conserve across kingdoms. For example, in some animal species such as dog and snake, hotspots are preferentially located around TSS and CpG islands [39, 40], whereas in some birds and plants they are associated with both TSS and TTS [19, 24, 30]. In plants, the fine scale recombination landscape has been characterized in a handful of species only, and the implicit view that plants have recombination hotspots mainly targeted to TSS and partly to TTS as in *A. thaliana* deserves further investigation. In addition, the simple observation of recombination gradient within genes can be misleading if the gene structure is not properly taken into account, as exemplified with gradients of GC content [36].

Here, we extended the characterization of fine-scale recombination landscapes in diverse flowering plants, specially focusing on genic regions. We assessed i) whether recombination hotspots are common in plants or whether some species lack them, such as *Drosophila* and *Caenorhabidtis*; ii) whether gene promoters and terminators are enriched in crossovers or whether hotspots can also be targeted outside genic regions in some species; and iii) how hotspot characteristics and gene structure (exons/introns) interact to shape the recombination gradient within genes. To do so, we constructed fine-scale recombination landscapes in eleven plant species, including nine new species (plus *A. thaliana* and rice), and we re-analyzed human data to compare with a species with PRDM9. We inferred fine-scale recombination rates from patterns of linkage disequilibrium in polymorphism data [41]. Owing to tiny distances between genetic markers (1-2 kb) and the large number of meioses observed in the genealogy of a population, variation in population-scaled recombination rates can be measured within genes and can be used to infer the precise hotspot locations [19, 22, 42, 43].

We found signature of hotspots in all eleven species but we uncovered a diversity of hotspot characteristics with preferential location in TSS (as assumed to be common) but also more frequent in TTS for some species, or without clear location around genic features. Despite this diversity, we propose that a simple model that takes gene structure (size, number of exons) and hotspot position into account can explain the variety of recombination gradients we observed within genes.

## Results

### LD-based recombination landscapes

To achieve fine-scale LD-based recombination maps at a gene scale, we gathered high-density polymorphism datasets in eleven flowering plant species (Table 1, Table S1). We identified species of interest, representing the diversity of plant genomes (small/large chromosomes, eudicots/monocots) and broad-scale recombination patterns (low/high genome-wide recombination rates), from a previous study [1]. We also used a human dataset [44] to compare the results with a species with hotspots targeted outside genic regions due to the PRDM9 mechanism. To ensure the comparability of the results we run the same pipeline on this human dataset instead of directly using published recombination maps. After filtering SNPs (minor allele frequency *>* 0.05, missing data per site *<* 0.1, bi-allelic SNPs, genotype quality score ≥ 30), we kept only datasets with at least 0.5 SNP/kb (range 0.6-9.7 SNP/kb, Table 1).

Fine-scale recombination landscapes have been estimated with LDhat [41] using a custom pipeline (https://github.com/ThomasBrazier/ldhat-recombination-pipeline.git v1.1). We checked for hidden population structure and sampled individuals within a single consistent genetic group with the highest polymorphism level, accordingly. For selfing species we considered haploid genome by sampling randomly one allele at the few heterozygote SNPs. The population size parameter (*θ_π_*) was estimated from the sampled population. The historical demography of the population was estimated with SMC++ [45] and used to generate a demography-aware look-up table with LDpop [46]. For computational limits, a maximum of forty diploid genomes per species (80 haploid genomes for selfing species, autosomes only) were sampled for LDhat analyses since the quality of estimates does not improve much over twenty individuals [47]. We checked the reliability of LD-based recombination landscapes by visual comparison (Fig. S1) between LD-based landscapes and broad-scale pedigree-based landscapes (Marey maps) of Brazier and Gĺemin [1]. We filtered all LD-based recombination maps to remove large genomic segments (*>* 100 kb) with a constant recombination rate (see the percentage masked in Table 1).

The mean genome-wide population-scaled recombination rates ranged from 0.44 to 12.34 *ρ*/kb across species (median range 0.03-4.6 *ρ*/kb). Most of the recombination rate estimates were lower than 10, yet the tails of the distribution were consistently skewed towards high values across species (Fig. S2). The ratio *ρ*/*θ* were globally in the same order of magnitude as in human (Table 1), except for *Camellia sinensis* (ratio ∼ 25.7). LD-based methods have been mostly evaluated on a range of parameters close to human, with a *ρ*/*θ* between 0.1 and 10 not affecting much the accuracy of estimates except for low mutation rates (*µ* = 10^−9^) [47].

**Table 1:**
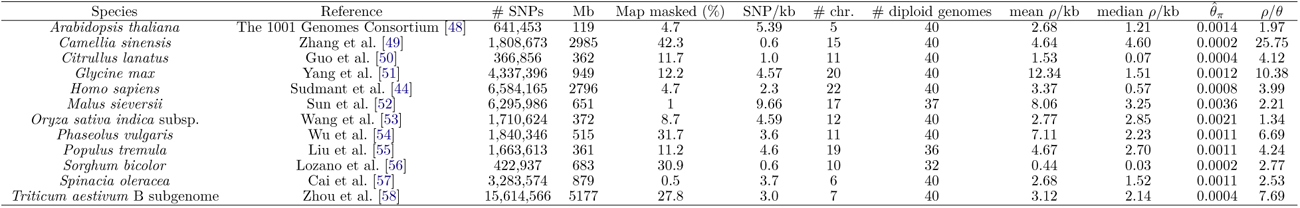
Summary statistics of twelve datasets (eleven plants and one human). Species, reference of the original polymorphism data, number of SNPs after filtering, genome length (Mb) and part of the LD map masked (%), SNP density (number of SNPs per kb), number of chromosomes of the genome, number of diploid genomes sampled, mean and median *ρ*/kb, estimated 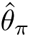 and ratio *ρ*/*θ*.

### Recombination hotspots are common but with different characteristics among species

Based on the LDhot statistical framework we inferred LD-based recombination hotspots in eleven plant species as well as in human in a standardized and comparable manner (Table 2, Fig. 1A). We applied two levels of filtering to the hotspot call set. As recombination hotspots are generally defined as narrow peaks of recombination over a short genomic range, we first filtered out all hotspots larger than 10 kb without making assumptions about their intensity (soft filtering). In order to improve our power to detect fine-scale patterns and associations with short targets (e.g. TSS, TTS), we did a harder filtering removing every hotspot larger than 10 kb and with an intensity lower than 4 and higher than 200 (putatively false hotspots due to variance in LDhat estimates). The hotspot intensity was measured as the peak rate divided by the background recombination rate (the mean background rate in a 50 kb window around the hotspot centre). The two levels of filtering drastically reduced the number of hotspots (Table 2). Hotspot filtering had the same impact on hotspot shape for all species. Hotspot size was reduced to approximately 6-8 kb and the hotspot intensity increased, as expected by the two criteria we chose (Fig. 1A).

The genome-wide hotspot density varied by a factor of 3.5 among species, independently of the filtering strategy (Table 2) and recombination rates varied over many orders of magnitude at a fine scale and were concentrated in a short fraction of the genome (Fig. 1B, Fig. S2). For most species, 80% of the total recombination was concentrated within approximately 20% of the genome. The species-specific patterns were roughly the same between plants and human. Our results for human, rice and *A. thaliana* were globally similar to state-of-the-art previous studies [19, 22, 42, 59], both for the shape of hotspots, the cumulative distribution function (Fig. 1A,B) and the number of hotspots (Table 2). Overall our results showed that recombination was globally concentrated in a short fraction of the genome across eleven plant species, suggesting that recombination hotspots may be a general feature in plants, similar to human.

**Table 2:** Number of hotspots (density of hotspots per Mb of the masked genome length). Statistics are given for the unfiltered, soft and hard filtering datasets. The number of hotspots already known in the literature is given at the end.

| Species | # hotspots | density/Mb | # (soft) | % (soft) | density/Mb (soft) | # (hard) | % (hard) | density/Mb (hard) | # (literature) | Reference |
| --- | --- | --- | --- | --- | --- | --- | --- | --- | --- | --- |
| *Arabidopsis thaliana* | 5,249 | 46.3 | 2,685 | 51.1 | 23.7 | 889 | 16.9 | 7.83 | 8,448 | [19] |
| *Camellia sinensis* | 41,843 | 24.3 | 14,098 | 33.7 | 8.18 | 5,299 | 12.7 | 3.08 |  |  |
| *Citrullus lanatus* | 9,068 | 28.3 | 4,511 | 49.7 | 14.1 | 2,726 | 19.0 | 5.4 |  |  |
| *Glycine max* | 69,499 | 83.4 | 66,287 | 95.4 | 79.5 | 23,467 | 33.8 | 28.2 |  |  |
| *Homo sapiens* | 104,573 | 39.3 | 57,952 | 55.4 | 21.7 | 16,952 | 16.2 | 6.4 | 25,000-50,000 | [22] |
| *Malus sieversii* | 26,286 | 43.8 | 13,814 | 48.8 | 21.4 | 4,054 | 14.3 | 6.3 |  |  |
| *Oryza sativa* | 12,443 | 36.6 | 6,134 | 49.3 | 18.0 | 3,251 | 26.1 | 9.5 | 14,125 | [30] |
| *Phaseolus vulgaris* | 13,550 | 38.5 | 7,573 | 55.9 | 21.5 | 2,886 | 21.3 | 8.2 |  |  |
| *Populus tremula* | 13,804 | 43.0 | 6,799 | 49.2 | 21.2 | 1,620 | 11.7 | 5.1 | 6,000 | [60] in *P. trichocarpa* |
| *Sorghum bicolor* | 12,350 | 26.1 | 6,324 | 51.2 | 13.4 | 2,859 | 23.1 | 6.0 |  |  |
| *Spinacia oleracea* | 75,376 | 86.2 | 73,431 | 97.4 | 84.0 | 30,413 | 40.4 | 34.8 |  |  |
| *Triticum aestivum* | 140,067 | 37.5 | 71,892 | 51.3 | 19.2 | 28,180 | 20.1 | 7.5 |  |  |

Despite qualitatively similar genomic sizes, hotspots varied in intensity among species (Fig. 1A, C). Three species exhibited particularly low levels of hotspot intensity compared to the background rate (*Citrullus lanatus*, *Malus sieversii* and *Phaseolus vulgaris*) while *Spinacia oleracea* and *Populus tremula* were the most intense plant species. Human presented an average pattern. Comparing the background recombination rate (± 50 kb) and the genome-wide rate, showed that hotspot detection was sometimes biased in favour of genomic regions recombining on average slightly less or more than the genome-wide average (see the difference between soft lines and the dashed line in Fig. 1A). For the unfiltered and soft filtered datasets, the bias was absent in three species and weak in eight other species (except *Citrullus lanatus*), suggesting that we were not strongly limited to detect hotspots in regions experiencing extremely high or low recombination rates. However, for the hard-filtered dataset, the detection of hotspots was particularly biased towards lowly recombining regions in many species (see the difference between the green and the dashed line), probably because intense hotspots are more easily detected on a low recombining background. In general, the soft-filtered dataset seemed relatively more reliable for the exhaustive search of hotspot location along the genome since hard filtering was biased towards low recombining regions in many species. Subsequently, we present results for the soft-filtered dataset.

**Figure 1:**
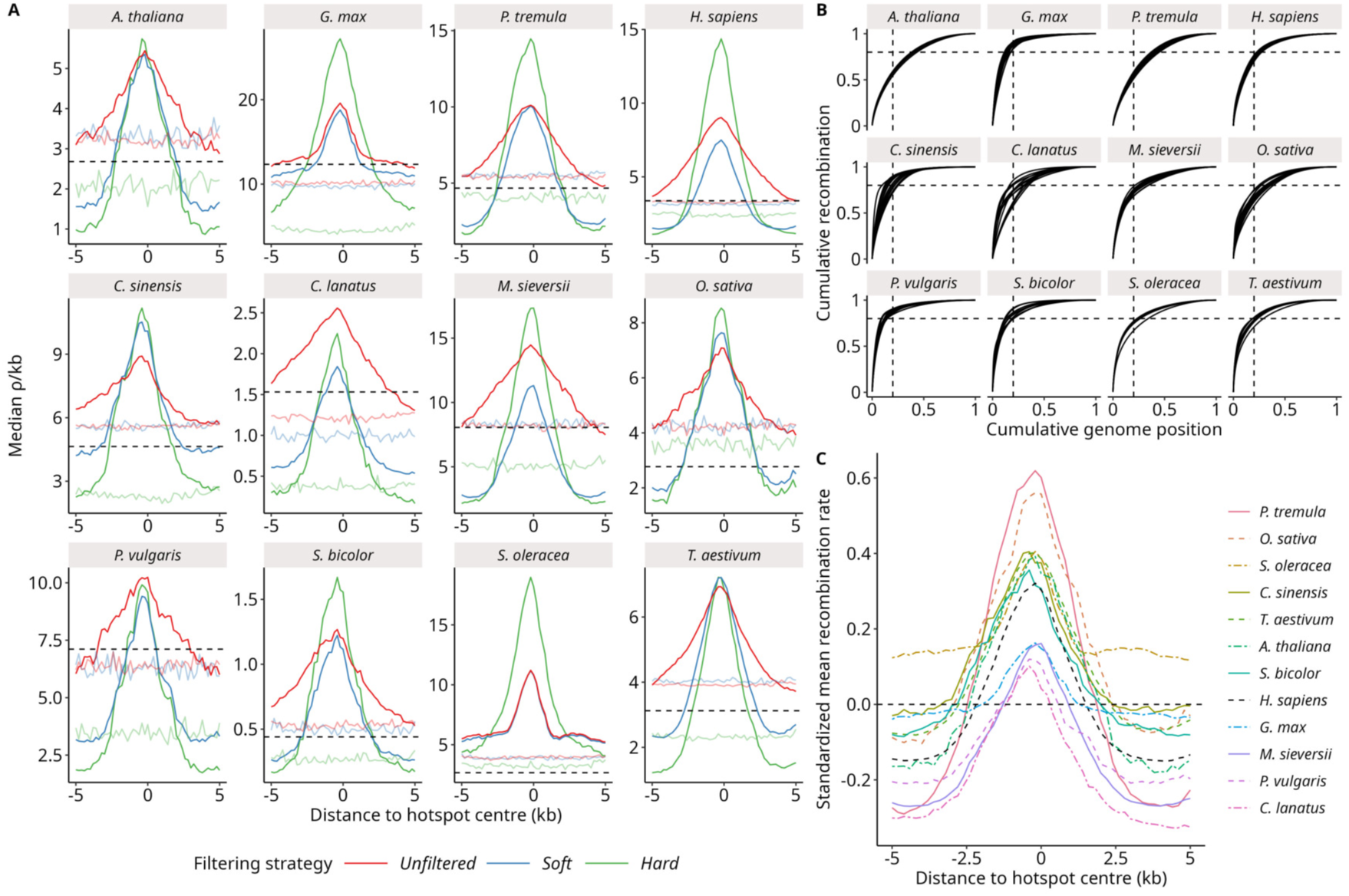
The distribution of recombination is heterogeneous at a fine scale and concentrated within recombination hotspots. (A) The recombination rate (*ρ*/kb) as a function of the genomic distance to the crossover hotspot centre (kb) with three different filtering strategies. Soft filtering removed hotspots larger than 10 kb. Hard filtering removed hotspots larger than 10 kb and hotspots with an intensity lower than 4 or higher than 200. Fully coloured lines are the recombination rates around hotspot centre (midpoint of the hotspot interval) and shaded lines are the control by randomly resampling hotspot intervals within a ± 50 kb neighboring region. The dashed horizontal line is the mean genome-wide recombination rate. (B) The cumulative ordered recombination fraction as a function of the cumulative genome fraction for each chromosomes and species. We represented dashed vertical lines at 20% of the genome and dashed horizontal lines at 80% of the total recombination. (C) Normalized (centered-reduced) recombination rates as a function of the distance to the hotspot centre for the soft filtered dataset. Species in the legend ordered by descending peak height. Human is in black.

### The fine scale distribution of recombination varies among species

As already observed in human, recombination rates were lower in genic than intergenic regions, except in *Populus tremula* and *Camellia sinensis*, even when flanking regions containing promoters and other cisregulatory elements are excluded (Fig. 2A). Here, we used the median instead of the mean to limit the effect of extreme values but using the mean recombination rates provided rather similar results with larger confidence intervals (Fig. S3A). Intergenic regions could be more prone to mapping errors, which could artificially inflate recombination rates. If it is a true signal, it suggests that recombination could also be directed towards intergenic regions, at least in some species. Despite lower recombination rate on average, genic regions were however enriched in recombination hotspots in eight over the eleven species: the number of hotspots overlapping a gene feature was significantly higher than expected (random expectation computed by 1,000 iterations of random shuffling of hotspot ranges) (Table 3, Table S2). Notably, it was also the case in human. In contrast, in *Glycine max*, *Spinacia oleracea*, *Triticum aestivum* the number of genic hotspots was less than expected just by chance.

Focusing on genic regions, recombination within Untranslated Regions (UTRs) and flanking regions was slightly higher than in coding regions (Fig. 2B, Fig. S3B). In most species, such as *Glycine max* and *Oryza sativa*, recombination rates were lower in UTRs than in flanking regions. However, in *Populus tremula* and *Camellia sinensis* recombination was only higher in the 3’ UTR and flanking region, with recombination being similar or higher in exons, introns than in 5’ part. We did not detect any clear pattern for differences in averaged recombination rates between exons and introns.

**Figure 2:**
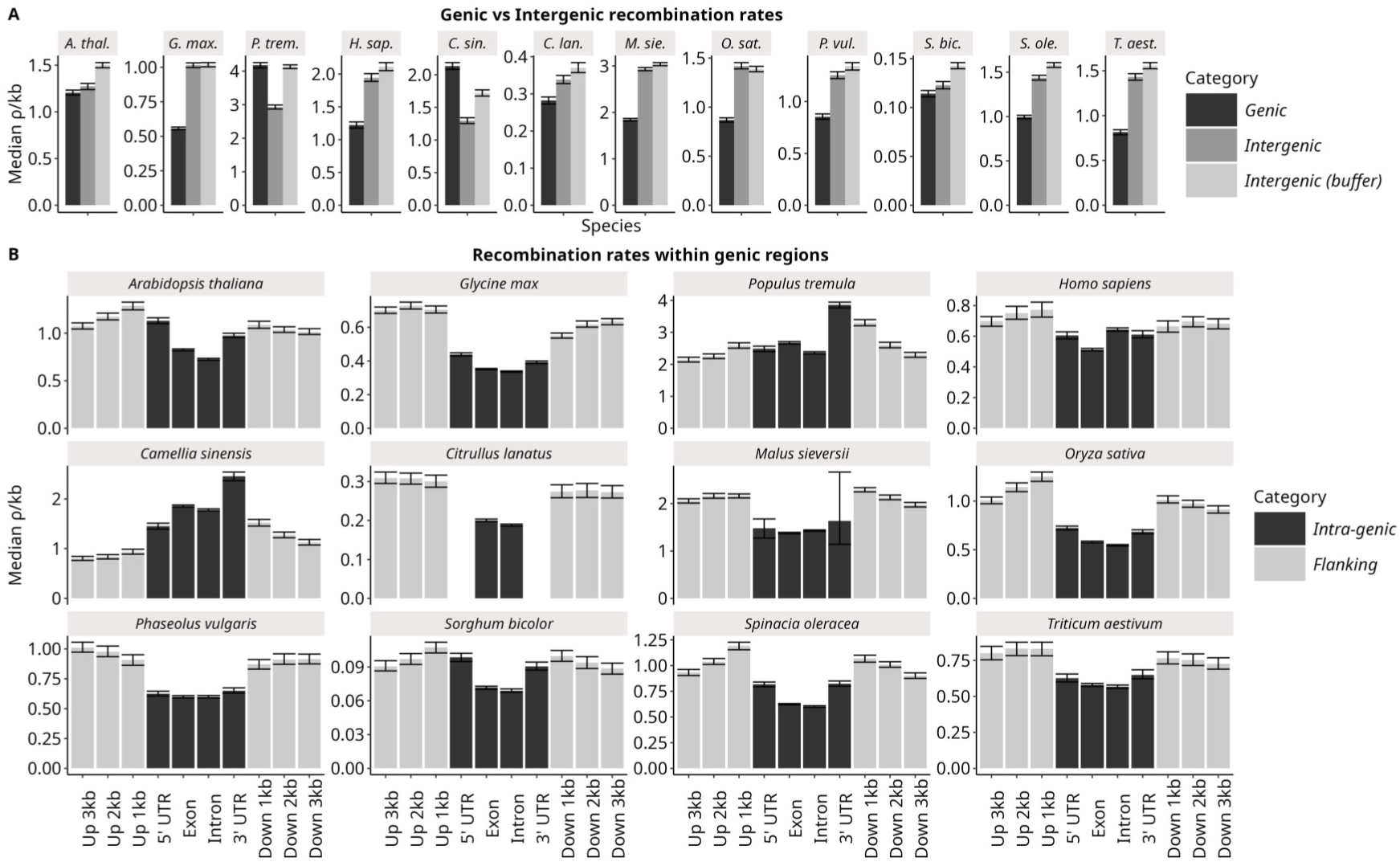
Variations in median recombination rates (*ρ*/kb) around and within genes. (A) Recombination rates vary between genic and intergenic regions. In order to remove an effect of 5’ and 3’ regulatory regions at the proximity of genes, buffered intergenic regions were defined by excluding 3 kb flanking regions upstream and downstream of genes. (B) Within genic regions, the recombination rate varies between genomic features as well as in 5’ upstream and 3’ downstream flanking regions. The median recombination rate and 95% confidence intervals were estimated by 1,000 bootstraps. Annotations of UTRs were not available for *Citrullus lanatus*.

**Table 3:** Number of genic and intergenic hotspots, i.e. hotspots overlapping or not a gene feature (soft filtered data). The expected number of genic hotspots was computed by 1,000 random shuffling of hotspots ranges.

| Species | # Intergenic hotspots | # Genic hotspots | # Expected genic (95% C.I.) |
| --- | --- | --- | --- |
| *Arabidopsis thaliana* | 215 | 2470 | 655 (614-700) |
| *Camellia sinensis* | 8449 | 5649 | 813 (760-867) |
| *Citrullus lanatus* | 2629 | 1882 | 687 (641-729) |
| *Glycine max* | 50076 | 16211 | 21144 (20917-21359) |
| *Homo sapiens* | 43122 | 14830 | 12894 (12703-13091) |
| *Malus sieversii* | 7414 | 6400 | 3101 (3011-3194) |
| *Oryza sativa* | 2263 | 3871 | 1147 (1090-1206) |
| *Phaseolus vulgaris* | 4308 | 3265 | 1348 (1287-1413) |
| *Populus tremula* | 1356 | 5443 | 1463 (1393-1528) |
| *Sorghum bicolor* | 3238 | 3086 | 759 (712-807) |
| *Spinacia oleracea* | 58161 | 15270 | 23883 (23645-24121) |
| *Triticum aestivum* | 68074 | 3818 | 11457 (11267-11631) |

Based on Figure 2, we chose three species (*Arabidopsis thaliana*, *Populus tremula*, *Glycine max*) illustrating the diversity of recombination patterns to characterise more precisely 5’-3’ genic recombination gradients. We found patterns similar to one of these three examples in other species (Fig. S4). In *A. thaliana*, the 5’ flanking region and UTR exhibited a rather large peak of recombination spanning about 1kb and recombination rate then steeply decreased from 5’ to 3’ just after the start codon. The 3’ end of the gene showed a small peak within the coding sequence just before the TTS but was much weaker than the 5’ end plateau. The 5’ peak was associated with a corresponding peak of hotspots density around TSS but we did not detect hotspot enrichment around TTS. This pattern was in agreement with previous results and considered as the standard pattern in plants [19]. *P. tremula* showed marked differences with the pattern observed in *A. thaliana*. We observed two narrow peaks inside the coding sequence, centred on the CDS start instead of the TTS and on the 3’ UTR instead of the TTS. Contrary to *A. thaliana* and *G. max* we also observed a higher peak in 3’ than in 5’, which both corresponded to an hotspot enrichment. Finally, in *G. max* we did not observed any peak but only a weak decrease towards the inside of the gene from both 5’ and 3’ ends, without hotpost enrichment (note also the different scales for the recombination rate).

Overall, these results clearly showed that the *A. thaliana* pattern is not universal in flowering plants and that some species, such as *G. max*, *P. vulgaris*, *C. lanatus* and possibly *T. aestivum* may not share the supposed ancestral system of targeting recombination towards promoter features or more generally gene flanking regions.

### Hotspot location and gene structure shaped gradients of recombination within genes

Previous results suggested that whatever the hotspot location, recombination gradients occurred inside genes. To go further we decomposed genic gradients per exons/intron ranks to remove spatial averaging effects among genes of different lengths and to evaluate the distribution of recombination within gene bodies. We kept only genes with less than 15 exons, as the sample size was too low above 14 exons (S3).

We first considered the average gradient by pooling all genes (see the black line in Fig. 4A). In *A. thaliana* and *G. max*, recombination decreased over the five to seven first exons and reached a minimal plateau with a slight re-increase at the end. In contrast, in *P. tremula*, after a decrease in 5’, recombination rates strongly re-increased from the middle of the gene to the 3’ end, yielding a U-shape gradient on average. We observed only slight differences in gradients among exons and introns (Fig. 4, Fig. S6).

**Figure 3:**
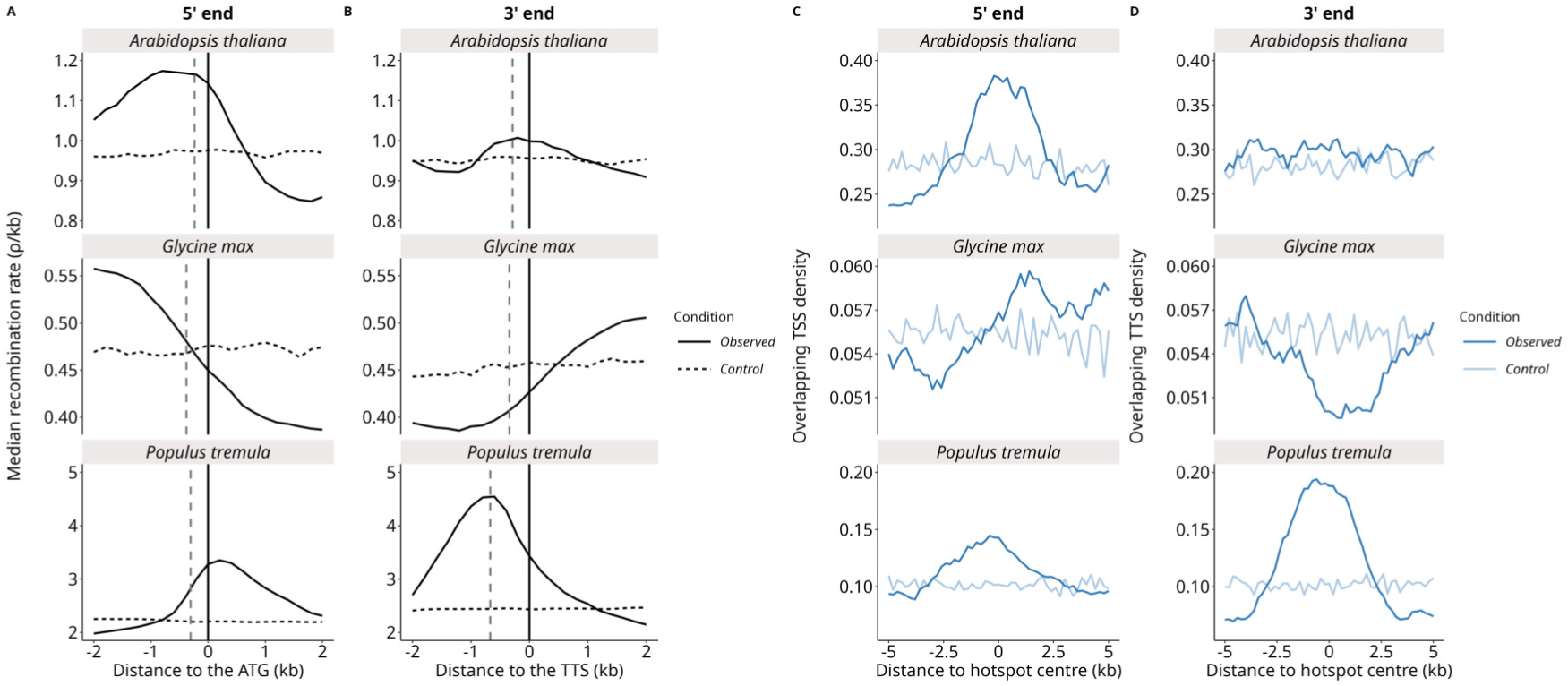
Recombination gradients along genes are mostly around Transcription Starting Sites (TSS) and Transcription Terminating Sites (TTS). (A, B) The median recombination rate (*ρ*/kb) was estimated in 200 bp windows as a function of the distance to the ATG start codon (start of first CDS) or the TTS and averaged among introns and exons. The vertical dashed lines are respectively the mean distance to the TSS (i.e. the 5’ UTR, panel A) and the mean 3’ UTR length (panel B). Random controls by randomly resampling intervals in the genome and re-scaled to the mean of the gradient. (C, D) The density of TSS and TTS as a function of the distance to hotspot centre (soft filtered hotspots). For smoothing, TSS and TTS were defined as regions of 1 kb centred on the gene start and end position respectively. Soft lines are a control by randomly resampling hotspot intervals in a ± 50 kb neighbouring region.

We then evaluated how gene structure (rank and number of exons) could shape differences in gradients among species. As such, gradients can also be observed by pooling genes according to their number of exons. We decomposed the averaged gradients across ranks into independent gradients for each exon-number class of genes (coloured lines in Fig. 4A). In all three species, shorter genes (genes with fewer exons) recombined more than longer genes on average (Fig. 4A, Fig. S7). However, it was not entirely driven by lower recombination rates in the middle of genes. The first exon(s) of shorter genes experienced higher recombination rates than the first exons of longer genes. Gradients per gene class were more or less parallel to the average gradient in *A. thaliana*, though a slight increase was detected in 3’ ends of longer genes (longer than seven exons). Gradients per gene class were weaker in *G. max* and the average gradient was mostly shaped by an excess of recombination in shorter genes (shorter than five exons). The structure of *P. tremula* gradients were more complex than the two previous patterns. The U-shape average gradient was produced by a combination of per gene class gradients increasing in 3’ and higher recombination rates in the first exons of shorter genes. In *P. tremula* gradients per gene class were strongly polarised towards the 3’ end. Importantly, the average gradient also depends of the proportion of each class of genes. As short genes are more frequent than long ones, this biases the average gradient towards the 5’ end.

**Figure 4:**
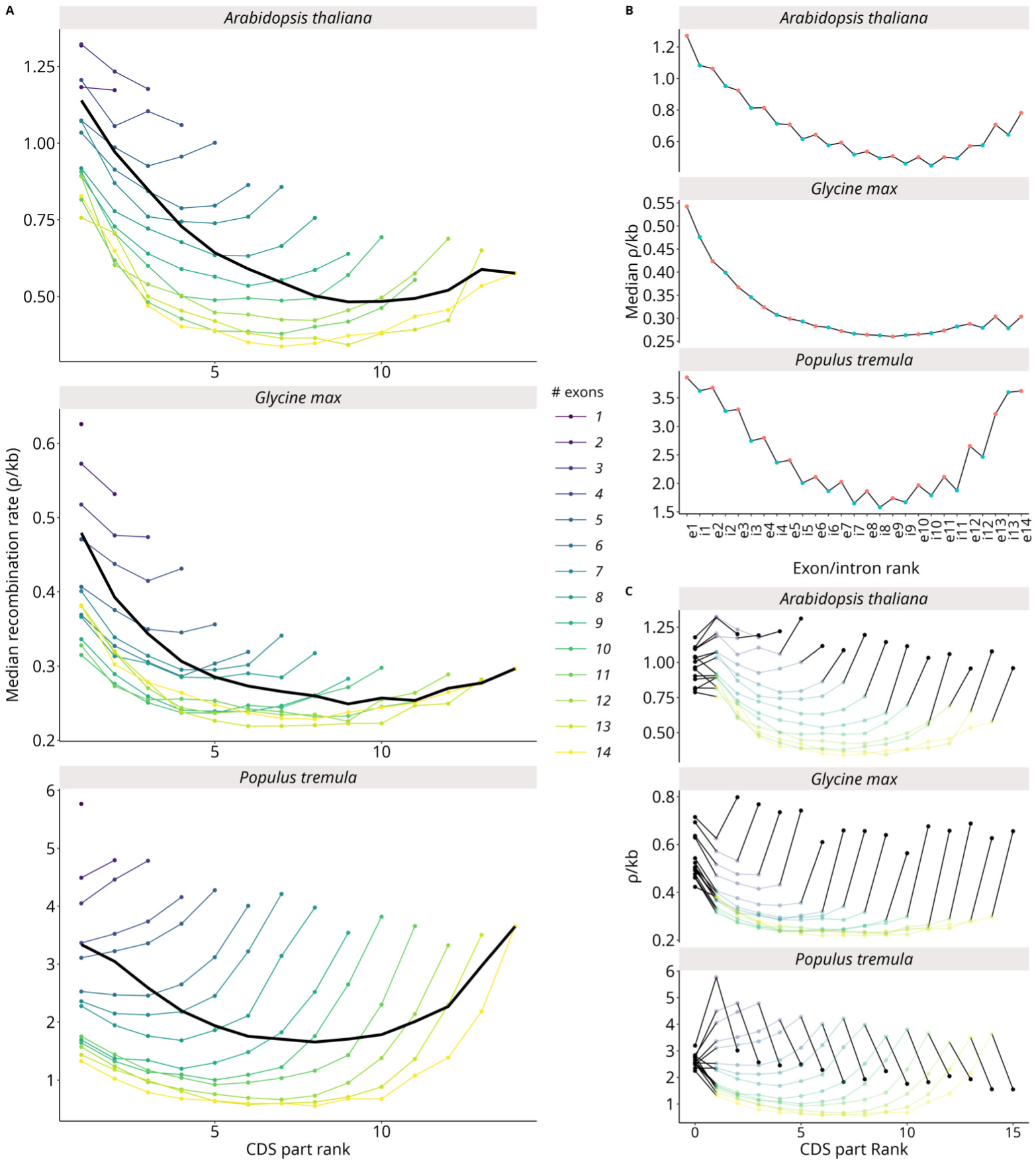
Gradients of recombination (median *ρ*/kb) along genes as a function of exon/intron rank. (A) Comparison of independent gradients as a function of exon rank (CDS part) with genes pooled by their number of exons. The black line is the average gradient (all genes pooled). (B) Average gradient of recombination rate as a function of exon of rank *i* (*e_i_*) or intron of rank *i* (*i_i_*). (C) Gradients in flanking regions representing the differences of recombination rates between genic regions and their flanking non-coding regions. On each side of the gene, the recombination rate of the 5’ and 3’ flanking regions (3 kb windows) is represented by a black dot and the difference between the gene part and its close flanking region is represented by a black line.

We checked if recombination rates at gene boundaries were correlated with the local recombination rate within flanking regions (Fig. 4C). In *A. thaliana* we noted a 5’ flanking recombination rate globally similar to the recombination rate of the first exon but elevated recombination rates in 3’ flanking regions compared to the last exon preceding them. In *Glycine max*, the differences with flanking regions was even stronger. The gradient within the coding sequence was weak compared to the increase in recombination rates in flanking regions. In *Populus tremula*, recombination rates at 5’ and 3’ flanking regions were similar among gene sizes and decoupled from variations within the transcript sequence.

### Recombination gradient inference is robust

LD-based recombination rate estimates are population-scaled and *ρ* is the product of the effective population size *N_e_* and the crossover rate *r* (*ρ* = 4*N_e_r*). As such *ρ* estimates are potentially biased if underlying variations in *N_e_* are not properly taken into account [61, 62]. Though we properly controlled for the genome-averaged effect of population structure and demography during the estimation of *ρ* (see material and methods for details), fine-scale variations in *N_e_* due to selection could also potentially impact LD-based estimates [63, 64]. To assess the robustness of our results to these potential biases we compared recombination gradients to fine-scale patterns of polymorphism (SNP density and genetic diversity *θ_π_*) pooled in the same manner (Fig. 5A, B). We observed a weak noisy gradient of SNP density in *Arabidopsis thaliana* but a clear absence of gradient in *Glycine max* and *Populus tremula* (Fig. 5A). The recombination gradients we observed are not likely byproducts of fine-scale variations in polymorphism. The ratio *ρ*/*θ_π_* followed a very similar gradients to *ρ*/kb. These results were strong evidence that recombination gradients were true gradients and not artifacts generated by underlying variations in selection (Fig. 5B).

To confirm that LD-based recombination rates were effectively correlated to estimates of crossover rates, we compared our results to the most resolved pedigree-based genetic map in the plant model *Arabidopsis thaliana* [65]. The two estimates were highly congruent despite different approaches and data. The LD-based *ρ*/kb mapped on 17,077 crossovers (pedigree-based) showed a peak of *ρ*/kb around the crossover centre (Fig. 5C). Conversely, the mean CO count around the LD-based hotspot centre showed that LD-based hotspots were enriched in crossovers (Fig. 5D). Despite a relatively low number of COs compared to the number of genes (17,077 COs vs. 25,942 genes), we were able to detect an increase in CO count around the TSS and TTS (Fig. 5E, F).

### The diversity of gradients across species can be explained by a single model

Applying the same analyses to the other plant species and to human we observed a diversity of patterns more or less similar to the three species we detailed above. Recombination patterns were mostly determined by differences in hotspot location and intensity. Average 5’-3’ recombination gradients were common across the eleven plant species and even in human (Fig. 6, black lines). The less defined gradient was in *Triticum aestivum* for which the power to detect gradients was limited (see methods for details). The level of LD was high in this species and the LD decay was very slow, even in genic regions, suggesting a lack of power to detect fine-scale patterns (Fig. S8). This diversity of recombination gradients was parallel to hotspot gradients (Fig. S5, Fig. S9). As a consequence, it seemed general among plants that shorter genes recombine more on average (Fig. 6, Fig. S7).

**Figure 5:**
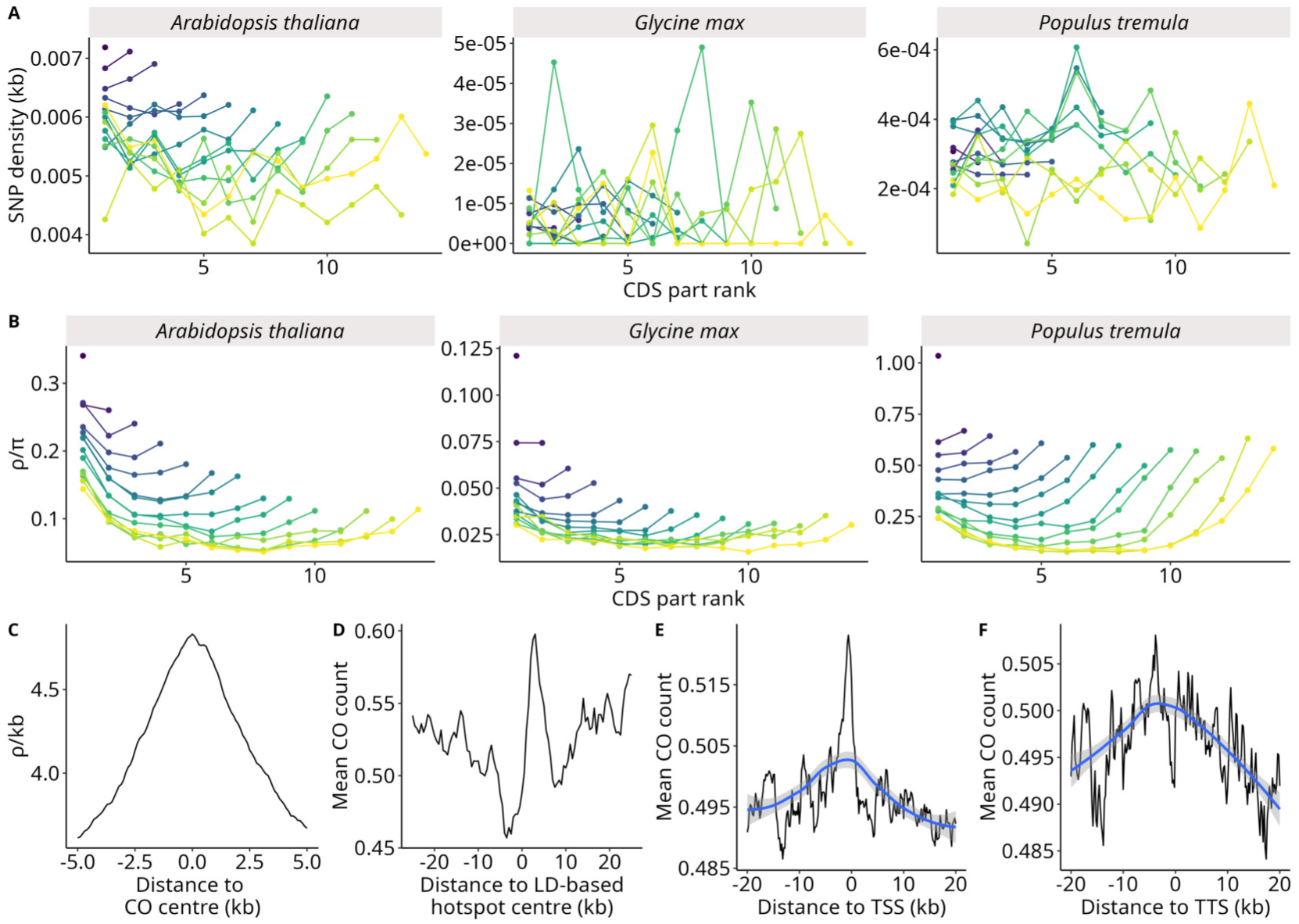
Recombination gradients are robust to variations in polymorphism levels and correlate with crossover rates estimated in Rowan et al. (2019). (A) Gradients of SNP density (SNP/kb) along genes as a function of CDS rank. SNP density was estimated for each CDS part and the average SNP density was calculated for each CDS part rank and gene size (number of exons). (B) Gradients of *ρ*/*θ_π_* along genes as a function of CDS rank. *θ_π_* was estimated per site and the average *θ_π_*/bp was calculated by the sum of *θ_π_*divided by the total sequence length of exons. *ρ*/*θ_π_* is *ρ*/kb divided by *θ_π_*/kb. (C) Peaks of LD-based recombination rates (*ρ*/kb) around crossover centre (± 5 kb) estimated in Rowan et al. (2019). (D) LD-based hotspots are enriched in crossovers. Mean crossover count around the hotspot centre (± 20 kb). Mean CO count measured by counting the number of crossovers overlapping 200 bp windows around the LD hotspot centre. (E,F) TSS and TTS are enriched in crossovers (± 20 kb). The blue line is loess regression with 95% C.I.

To better understand this diversity of patterns, we formalized a simple model based on the Choi and Henderson conceptual model (2015). We assumed that recombination exponentially decreased within gene from both ends, starting with recombination rates in 5’ and 3’ flanking regions, *r*_5_ and *r*_3_, depending on hotspots location and intensity (see equations 1-3 in Material and Methods). Under this mechanistic view, exon and intron lengths, which vary with gene structure (Fig. S10) should play a role and took them explicitly into account. Because we focused on within gene gradients, this model corresponds either to hotspots located in 5’ and 3’ regions with different proportion and/or intensity, such as in *A. thaliana* and *P. tremula*, but also if hotspots are targeted away from genic regions, with no recombination peak in 5’ and 3’ but a simple decrease in recombination rate within genes, as in *G. max*. We fitted this model to the detailed gradients shown in Fig. 6. This simple model fitted the data very well (*r*^2^ from 0.64 to 0.96, Table 4, Fig. 7B, C, Fig. S12) and the ratio *r*_5_/*r*_3_ was congruent with the observed gradients.

**Figure 6:**
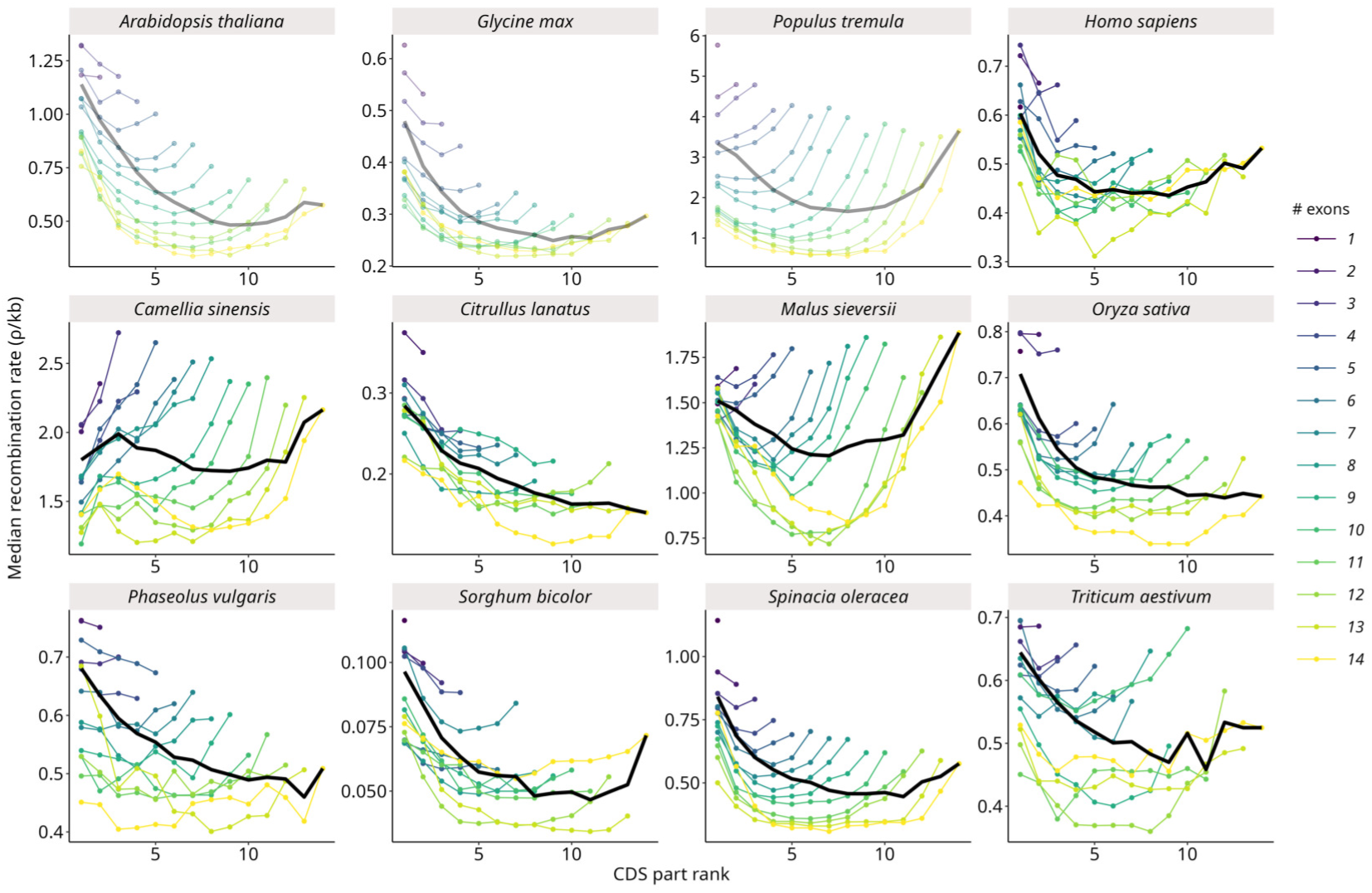
Diversity of recombination gradients across plant species. The recombination rate (median *ρ*/kb) was estimated in exons (CDS part) as a function of their rank and genes were grouped by their number of exons. The black line is the average gradient (all genes pooled). The three reference species already shown above are transparent. Only protein-coding genes were used.

**Table 4:** Fit of the model and parameter estimates for eleven species. The basal recombination rate *r*_0_ (standard error), the 5’ recombination rate *r*_5_ (standard error), the 3’ recombination rate *r*_3_ (standard error), the ratio *r*_5_/*r*_3_, the inverse of the tract length *a* (standard error), the tract length in bp, and the *R*^2^ of the Non-Linear Least Squares fitting procedure. The standard errors of the predictions were obtained with the ‘nls’ R function.

| Species | $r_0$ | $r_0$ (s.e.) | $r_5$ | $r_5$ (s.e.) | $r_3$ | $r_3$ (s.e.) | ratio $r_5/r_3$ | $a$ | $a$ (s.e.) | Tract length (bp) | $R^2$ |
| --- | --- | --- | --- | --- | --- | --- | --- | --- | --- | --- | --- |
| *Arabidopsis thaliana* | -0.74 | 0.09 | 1.43 | 0.05 | 1.17 | 0.05 | 1.23 | 0.000505 | 0.00003 | 1,982 | 0.95 |
| *Camellia sinensis* | 0.49 | 0.24 | 0.69 | 0.11 | 1.59 | 0.16 | 0.44 | 0.000128 | 0.000023 | 7,817 | 0.76 |
| *Citrullus lanatus* | -3.58 | 1.29 | 2.35 | 0.7 | 1.62 | 0.59 | 1.45 | 0.000042 | 0.000011 | 23,950 | 0.72 |
| *Glycine max* | -0.01 | 0.04 | 0.35 | 0.02 | 0.28 | 0.02 | 1.26 | 0.000302 | 0.000031 | 3,315 | 0.84 |
| *Homo sapiens* | 0.36 | 0.02 | 0.23 | 0.02 | 0.15 | 0.02 | 1.56 | 0.000067 | 0.000012 | 14,962 | 0.71 |
| *Malus sieversii* | 0.77 | 0.07 | 0.66 | 0.06 | 0.99 | 0.06 | 0.67 | 0.000629 | 0.000076 | 1,590 | 0.78 |
| *Oryza sativa* | -0.07 | 0.06 | 0.56 | 0.04 | 0.48 | 0.03 | 1.15 | 0.000294 | 0.000028 | 3,399 | 0.86 |
| *Phaseolus vulgaris* | -0.08 | 0.09 | 0.48 | 0.05 | 0.46 | 0.05 | 1.04 | 0.000175 | 0.000026 | 5,712 | 0.84 |
| *Populus tremula* | -1.77 | 0.23 | 3.4 | 0.14 | 5.38 | 0.15 | 0.63 | 0.000442 | 0.000025 | 2,263 | 0.96 |
| *Sorghum bicolor* | -0.02 | 0.02 | 0.09 | 0.01 | 0.06 | 0.01 | 1.44 | 0.00028 | 0.000047 | 3,576 | 0.69 |
| *Spinacia oleracea* | -0.05 | 0.03 | 0.65 | 0.02 | 0.59 | 0.02 | 1.1 | 0.000239 | 0.000013 | 4,180 | 0.95 |
| *Triticum aestivum* | -0.04 | 0.13 | 0.44 | 0.07 | 0.43 | 0.07 | 1.01 | 0.000186 | 0.000041 | 5,365 | 0.64 |

**Figure 7:**
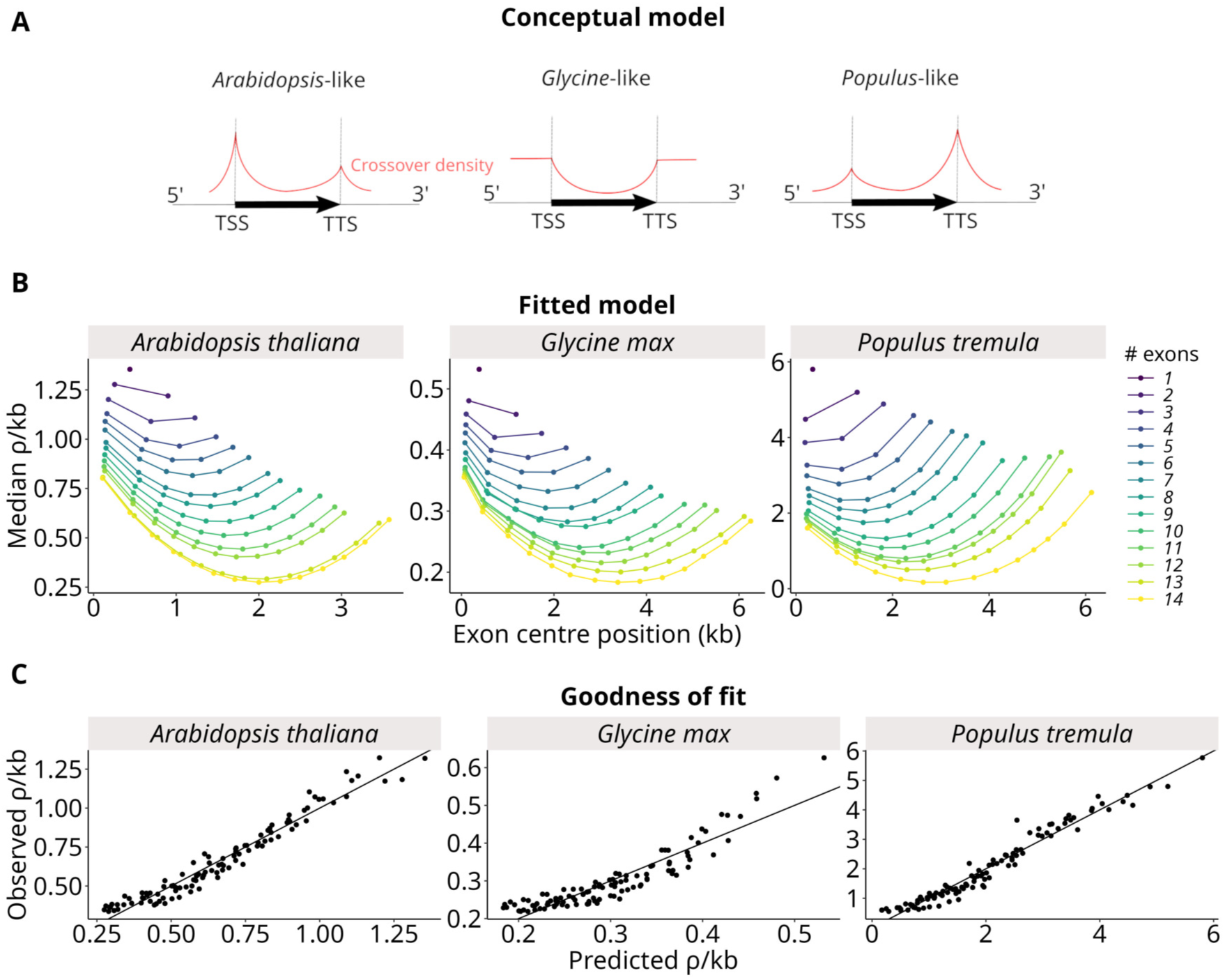
Fit of the model to recombination gradients. (A) Conceptual model of recombination gradients adapted from Choi and Henderson (2015). Recombination rates exponentially decay around the TSS and TTS. (B) Recombination rates predicted by the model, as described in equations 1-3. Model fitted with Non-Linear Least Squares and the sum of square differences between expected and observed recombination rates as a loss function to minimize. (C) Goodness of fit of the model, with observed values as a function of predicted values.

## Discussion

We characterized recombination patterns in genic regions in eleven plant species, including nine species not described before. We detected recombination hotspots in all species and we generalised the previously observed pattern that recombination hotspots are organised around the boundaries of coding sequences, both at the 5’ and 3’ ends of genes. However, we also uncovered more variations than initially envisaged from only a few model species, including some species without clear targeting of recombination hotspots around genic regions. We also leveraged high-resolution recombination maps at the exon scale to characterize detailed genic recombination gradients and we proposed a single model that can explain the diversity of observed patterns through the interaction between hotspot location and gene structure.

### Recombination varies around and within coding sequences

At the gene scale, the location of recombination hotspots was often concentrated in gene 5’ and 3’ ends, as previously known for a few plants only [19, 28–30, 66]. We observed that recombination can be targeted towards diverse domains both in 5’ and 3’ ends (UTRs, TSS/TTS, promoters, flanking) and was not systematically associated with promoters as previously found [28, 29, 67]. Despite divergent patterns, it is clear that the genic region is finely structured according to the different genomic domains composing it.

Within 5’ and 3’ ends enriched in COs in *A. thaliana*, chromatin accessibility redirects meiotic recombination towards gene promoters and terminators [19], and the same mechanism also likely occurs in other species [3]. In plants as in yeast, double-strand breaks (DSB) hotspots, the precursors of COs, are found both in TSS and TTS where nucleosome occupancy is reduced whereas COs form close to H2A.Z and H3K4me3 histone marks which are epigenetic modifications of chromatin packaging also involved in the regulation of gene expression [19, 68, 69]. Low methylation is also usually associated with CO hotspots [19]. Both TSS and TTS are hypomethylated in *A. thaliana*, *G. max* and *P. tremula* [70–72], which fits well with our results. These findings are globally in agreement with the tethered-loop/axis model, which postulates that DSBs form globally in chromatin loops bound to the chromosome axis whereas COs occur only in nucleosome-depleted regions that are susceptible to being tethered to the central chromosome axis by recombination-promoting factors [4, 73]. Indeed, gene promoters and terminators are located in chromatin loops and COs most probably concentrate here due to their highly conserved chromatin state and DNA accessibility [4, 74]. However, *P. tremula* and *C. sinensis* do not conform exactly to this canonical model as CO hotspots preferentially occur inside rather than outside coding sequences, as also seen in *Mimulus guttatus* [29]. In *PRDM* 9^−^*^/^*^−^ mice, the centre of PRDM9-independent hotspots is on the +1 nucleosome positioned downstream of the TSS [18]. For these species, the detailed description of chromatin and histone patterns during meiosis (especially H2A.Z and H3K4me3), as well as nucleosome maps, could be useful to better understand this non-canonical pattern [3, 4].

We also identified species, *G. max*, *P. vulgaris*, *C. lanatus* and *T. aestivum* (Fig.S4, Fig. S5), where we did not identify a clear signature of hotspot location around 5’ and/or 3’ regions, and some species with hotposts possibly targeting away from genic regions *G. max*, *S. olearacea*, and possibly *T. aestivum*, which showed a significant deficit in hotspot in genic regions (Table 2). Interestingly, *P. vulgaris* and *C. lanatus* showed an excess of hotspot in genic regions without clear 5’/3’ pattern whereas *S. olearacea* showed a deficit of hotspots in genic regions but clear peaks both around TSS and TTS. If it is a true signal, it would suggest the existence of additional mechanisms, different from the supposed ancestral mechanism, which can direct recombination towards other targets than promoter-like sequences. In addition, also in contrast with previous expectations, we found a signature of 5’ hotspots with a weak 5’-3’ recombination gradient in human, despite the PRDM9 mechanism. This reinforces the recent findings in some animals that several competitive mechanisms could be at play in locating recombination hotpots within a genome [38, 39].

### Genic recombination gradients are shaped by hotspot location and gene structure

The specific location of hotspots can generate recombination gradient within genic regions, which can be used as a signature to distinguish promoter-driven and PRDM9-driven mechanisms [24, 39]. However, in previous studies, only average gradients were considered. We showed that it is key to decompose recombination gradients as a function of exon rank and gene length (in exon number here) to obtain a proper characterization of recombination patterns and better understand the underlying mechanism. Typically, because short genes are more frequent than longer ones, the average gradient is biased towards the 5’ end, such that J-shaped patterns can appear as U-shaped, as in *P. tremula* or almost flat pattern can appear as a standard 5’-3’ gradient as in *P. vularis*.

The intensity and location of hotspots are globally responsible for the polarity of recombination within genes. Hotspots overlapping 5’ and 3’ ends of genes have a consistent influence on the average recombination rate. More importantly, recombination hotspots seem similar in size but their intensities vary between species (Fig. 1, Fig. S3). Hotspot position also shapes the recombination gradient towards the 5’ or 3’ end. Though the TSS is usually the preferential location of CO hotspots, the increasing gradient towards the 3’ end in *P. tremula*, *M. sieversii* and *C. sinensis* is associated with the location of a significant fraction of recombination hotspots around the TTS [20]. In other species, such as *G. max*, the intensity of hotspots compared to the genome-wide average recombination rate is rather low and TSS and TTS do not seem to be enriched in hotspots in *G. max*, which can explain why gradients are less pronounced in such species. Our modelling approach showed that these various patterns can be explained under a single framework taking hotspot location (around or away 5’ and 3’ flanking regions) and gene length and structure into account. This model fits all observed pattern well, even novel patterns undescribed until now, and provides a quantitative measure of the relative asymmetry between 5’ and 3’ hotspots. Note that in human, introns are extremely large compared to plants (Fig. S9B) and could explain why the 5’ gradient is steep and mostly concentrated in the first exon.

Gene length not only affects recombination in the middle of genes but it also lowers recombination for all exon positions. As the proportion of hotspots is invariant with gene length, the same amount of recombination is automatically spread over more exons, making average *ρ*/kb more or less negatively proportional to gene length. The phenomenological model we propose following Choi and Henderson (2015) can explain this observation simply because shorter genes have a closer influence of hotspots on both sides than longer genes (Fig. 7). A direct consequence is that shorter genes recombine more than longer genes on average, as we observed in nine out of twelve species.

Finally, our results support the prediction that the similar GC gradients shaped by exon rank and gene class observed in plants can be generated by the underlying recombination gradients we characterized through the effect of GC-biased gene conversion [36]. In *A. thaliana* and rice, recombination gradients per rank and gene class match GC gradients relatively well (compare our Fig. 6 with Fig. 5 in Ressayre et al. [36]).

### LD-based estimates are reliable to detect gradients

We addressed the limitations and reliability of LD-based estimates of recombination rates. We carefully controlled for population structure, mating system and demography during the estimation procedure, following the most recent developments and benchmarks in the literature [46, 47]. A recent simulation-based study showed that LDhat was the most accurate method to estimate fine-scale estimates of recombination rates from polymorphism data in presence of hotspots [75]. LDhat was even more accurate when it was combined with LDpop [46]. In *Arabidopsis thaliana* LD-based estimates correlated well with the Rowan et al. (2019) pedigree-based CO map. The power to detect a gradient was not affected by the mating system or data quality (SNP density). The highly selfing *Arabidopsis thaliana* is one of the most resolved gradients. We observed a clear gradient in *Camellia sinensis* despite quite low SNP density (0.6 SNP/kb) while the gradient was noisy in *Triticum aestivum* with five times more SNPs (3 SNP/kb). We also checked that recombination gradients were not biased by local patterns of polymorphism. Altogether, the extremely fine resolution we achieved in our analyses proves the good accuracy and reliability of our pipeline, which is freely available and easy to use (https://github.com/ThomasBrazier/ldhat-recombination-pipeline.git).

## Conclusion

It was known that many organisms, including plants, had recombination hotspots [9, 76, 77], sometimes inducing recombination gradients in genic regions. Our results confirm that it is likely a rather general feature of plant genome but we also discovered an unexpected diversity of patterns. Indirect approaches like ours are important to detect unseen patterns and open new questions for future research, such as the mechanistic bases behind the 3’-driven gradient in *Populus tremula* and *Camellia sinensis*.

We also provided a new model taking gene structure into account to unravel the complexity of these recombination gradients that likely emerged from both the location of recombination hotspots and the distribution of genes per class of exon numbers. Our modelling approach allowed us to highlight the relative importance of hotspots in 5’ and 3’ for the shape and intensity of recombination gradients.

We suggest that a detailed characterisation of recombination and its known covariates (e.g. methylation, chromatin state) at such a fine scale will be helpful to better understand the evolution of recombination hotspots themselves [3, 78], patterns of base composition in genic regions [35] or the evolution of intron structure [79]. Moreover, the evolutionary significance of these recombination gradients in genic regions may have been underappreciated so far, as they scale with effective population size over thousands of generations and might have a deep impact on coding sequences through complex interactions with linked selection or gBGC [35, 80].

## Materials and Methods

To produce fine-scale recombination maps we analyzed previously published polymorphism datasets in eleven plant species and human (Table 1) starting from VCF files. We identified datasets by literature search with the keywords “resequencing”, “high-density SNP”, “polymorphism data”, “variant database”, “Whole Genome Sequencing” or “WGS” in association with “plant”, “angiosperms” and a set of plant species names. We also searched directly for public genomic databases (e.g. 1001 genomes). We used a previous study of 57 plant species to identify species with interesting and contrasting characteristics, such as monocots/eudicots, small/large genomes, low/high recombination rates, and homogeneous/heterogeneous recombination landscapes [1]. Before downstream analyses, we kept only bi-allelic SNPs and filtered them with the same quality criteria for all datasets (minor allele frequency *>* 0.05, *<* 10% of missing data per site, quality score ≥ 30 when the information was available). When the quality score was not available in the published VCF, we checked in the original study that the dataset we re-used was already filtered by a similar quality score.

Genome assemblies and corresponding annotations (GFF files) on which SNPs were originally mapped were retrieved from public databases, mostly NCBI (Table S1). GFFs with gene, mRNA, CDS and exon features were parsed using custom Python scripts to infer introns, 5’ and 3’ UTRs, flanking sequences (1-3 kb upstream/downstream, in consecutive windows of 1 kb) and intergenic sequences according to the GFF3 specifications. Scripts are freely available as a Python package (https://github.com/ThomasBrazier/PiSlice. git v0.2). Only protein-coding genes were retained when this information was annotated. For datasets without information about gene biotypes, we considered all genes as protein-coding. To avoid low sample size for longer genes, we removed genes with more than 14 exons as in Ressayre et al. (2015). When more than one splicing variant has been annotated, the splicing variant with the longest CDS sequence was retained, as it is often also the most expressed sequence. We considered only the CDS part of exons.

All statistical analyses were performed with R version 4.3.3 [81]. Genomic coordinates were manipulated with the R package GenomicRanges [82].

We developed a method for estimating robust LD-based recombination rates and implemented it in a custom pipeline performing data processing, SNP filtering, statistical phasing, controlling for population structure and demography, dealing with selfing and highly homozygous species, estimating recombination rates with LDhat 2.2 [41, 83] and inferring hotspots with LDhot [42]. This pipeline was optimized to run large datasets with millions of SNPs. The Snakemake pipeline is freely available (https://github.com/ThomasBrazier/ldhat-recombination-pipeline.git v1.1) and the method is presented in detail below.

### Population sampling

Population-scaled recombination rates and hotspot detection are based on the assumption of a population without genetic structure and absence of gene flow from other populations [61, 62]. For each dataset, we inferred population structure and sampled one single genetic population as homogeneous as possible and well distinct from other possible genetic clusters. As much as possible, we tried to sample a single large population (*>* 40 diploid individuals or 80 haploid genomes for selfing species) of wild/landrace individuals with the highest possible polymorphism level. Results and metadata of the original publications were used for a pre-sampling. The genetic structure within the pre-sampled population was assessed with FastSTRUCTURE for K = 1-7 [84]. FastSTRUCTURE was run on a random subset of 100,000 SNPs for computation issues. The best model (best value of K) was inferred by FastSTRUCTURE’s algorithm for multiple choices of K and validated by the visual assessment of admixture plots. If a population substructure was positively assessed, individuals were clustered into different genetic populations according to their mean admixture proportion. Clusters with higher proportions of missing data and low polymorphism (heterozygosity and *θ*^^^*_π_*) were discarded and we kept individuals in the largest remaining genetic cluster.

### Population-scaled recombination rates

After population sampling, the VCF was phased with Shapeit2, with a phasing window size of 2 Mb [85]. Selfing species were treated as haploid individuals, by randomly sampling only one phased haplotype per individual. When the sample size was sufficient, at most 40 diploid individuals (or 80 haploid sequences for selfing species) were randomly subset for LDhat 2.2 for computational issues (see Table S1 for sampling sizes).

The population-scaled recombination rates *ρ*/kb were estimated with LDhat 2.2 [41, 83]. For the estimate of genetic diversity required by ‘LDhat’ we used the genome averaged 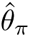 estimated with ‘VCFtools’ on the population dataset [86]. The per-base mutation rate was set to 1.28 ∗ 10^−8^ for all species. Previously to ‘LDhat’, we estimated the demography of the population per chromosome with ‘SMC++’ [45] with the default parameters, including eight epochs, and a missing cutoff for runs of homozygosity adjusted per chromosome (see Table S1 for parameters). We used ‘LDpop’ to generate a complete demography-aware look-up table with the *θ* estimate and the fast approximate method [46]. The maximum value of *ρ* was set to 100, as suggested in the ‘LDhat’ manual. Recombination rates between pairwise adjacent SNPs were estimated with the ‘LDhat interval’ program, using different values of block penalty (bpen = 5, 15, 25). Since the quality of recombination landscapes was similar between different block penalties, despite a smoothing effect for higher values, we kept estimates with the smaller block penalty (bpen = 5). For computational efficiency of the ‘LDhat interval’ part in larger datasets, the dataset was split into intervals of 2,000 SNPs (50 SNPs overlapping at each end), run in parallel and merged at the end of ‘LDhat interval’. We ran the MCMC algorithm for 10,000,000 iterations, sampling every 5,000 iterations, with a burn-in of 1,000,000 first iterations. In a few species we adjusted these hyper-parameters to reach convergence (Table S4). The convergence of MCMC sampling chains was checked with ‘LDhat’ diagnostic plots. For species with an already published Marey map in Brazier and Gĺemin (2022), the visual congruence between the LD map and the Marey map was used as a qualitative validation of our estimates.

### Hotspot detection

Recombination hotspots were inferred with ‘LDhot’ using default settings [42]. We tested the significance of putative hotspots in 3 kb windows, with a step size of 1 kb along the genome. The background window was ±50 kb around the hotspot centre. The hotspots inferred were then summarised using the ‘LDhot’ summary program with a significance cutoff of 0.001 for calling a hotspot and 0.01 for merging adjacent hotspots. In order to control for hotspot quality and reduce false positive hotspots, we compared the raw hotspot dataset to filtered datasets after soft and hard filtering strategies. As hotspots are generally defined as narrow genomic regions, the soft filtering approach removed every hotspot larger than 10 kb. The intensity of a hotspot was estimated by dividing the recombination peak rate by the background recombination rate (±50 kb around the hotspot centre). Low-intensity hotspots were considered to be most likely false positive calls. In addition, we considered that extreme values of hotspot intensity were most probably artefactual due to high variance in the LDhat recombination map caused by assembly and/or SNP calling errors. The hard filtering removed hotspots larger than 10 kb or with an intensity lower than 4 or higher than 200.

### Recombination rates and hotspots in genomic features

We calculated the median and average *ρ*/kb for each genomic feature with the median and the weighted arithmetic mean of the *ρ*/kb estimates intersecting this genomic interval (R ‘median’ and ‘weighted.mean’ function). The weights were the number of nucleotides spanned by each interval. The number of hotspots overlapping a given genomic feature was counted with ‘countOverlap’ and ‘findOverlap’ from the GenomicRanges R package [82]. The 95% confidence intervals of the mean and the median of *ρ*/kb within genomic features were estimated by bootstrap (1,000 iterations).

### Gradients of recombination

To assess a putative gradient of recombination at the 5’ end of genes, we estimated LD-based recombination rates (*ρ*/kb) in consecutive intervals (200 bp) along a region ±5 kb around the TSS, ATG codon (start of the coding sequence) and TTS. Annotations of start codons are generally more consistent and reliable than Transcription Starting Sites (TSS) among genomes that have been annotated using different methodologies. Intervals overlapping another gene were discarded. We also separated the average gradient from exon/intron-specific gradients. Each interval in a gene was marked as exon-specific (intron-specific, respectively) only if an exon (intron) overlapped at least 70% of the interval. A random control of *ρ*/kb was calculated by resampling the same intervals at a random position in the genome.

We also produced gradients of recombination by estimating recombination rates as a function of exon/intron ordinal rank within genes as in Ressayre et al. [36]. We computed the mean, weighted mean and median *ρ*/kb for each exon (CDS part) and intron. The recombination rate of exons/introns was averaged by exon/intron ordinal rank for the average gradient, and by rank and gene size (number of exons) for detailed gradients. As annotation quality and sample size declined for longer genes, we removed all genes longer than 14 exons or 10 kb.

We evaluated the performance of our method to produce gradients by testing whether it had the power to detect a true gradient of recombination or the capacity to reject an artifactual gradient. Firstly, we assessed whether the gradient observed could be an artefact. LDhat recombination rates are estimated within intervals between adjacent SNPs. As a consequence, a lack of SNPs at the beginning of genes where recombination hotspots are supposed to be located could reduce our resolution to accurately position hotspots in this region, thus leading to an artificial leakage of high *ρ*/kb values at a larger scale than the true biological size of the hotspot. To evaluate this, we re-calculated the gradient of *ρ*/kb as a function of the rank, but this time we kept only *ρ*/kb estimates beginning in the genomic interval (exon/intron). Thus, we were sure to measure only the actual recombination within the interval. We observed the same patterns using this restricted set of *ρ* values confirming that the observed gradients were a true biological signal (Fig. S13A, B). Secondly, we assessed the power to detect a true gradient, if there was one, for a given empirical dataset. For each gene, the ranks of exons were sorted by descending *ρ*/kb values, to simulate the maximal negative 5’-3’ gradient that could be observed from empirical observations, if there is one. Ten out of eleven datasets had the necessary power to detect true gradients of recombination, except for *Triticum aestivum* lacking resolution (Fig. S13B).

### Fitting of the hotspot model

We fitted a phenomenological model to recombination gradients explicitly taking into account exon positions (starting at *x_s_*and ending at *x_e_*), exon length (*L_exon_*= *x_e_* − *x_s_*), and gene length (*L_tot_*). We considered that recombination within gene was generated by two exponentially decreasing gradients starting from 5’ and 3’ ends, respectively, plus a relative basal rate *r*_0_, which can be negative. We assumed a same decreasing rate, *a*, for each end. If hotspots were centred exactly at the beginning and at the end of genes, 1*/a* could be interpreted as the length of the recombination tract. However, depending of the species, recombination can start to decrease before of just after gene boundaries. Accordingly, recombination rates in 5’ (*r*_5_) and 3’ (*r*_3_) do not necessarily correspond to hotspot rates. More precisely, *r*_5_ = *r*_5_*_,max_e*^−^*^ad^*^5^, where *r*_5_*_,max_* is the recombination rate at the centre of the hotspot and *d*_5_ is the distance between the hotspot centre and the start of the gene (similar interpretation for *r*_3_). Unfortunately, *r*_5_ and *d*_5_ (respectively *r*_3_ and *d*_3_) are not identifiable. For each exon, the predicted recombination rate is thus given by:

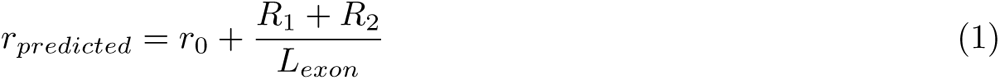

with

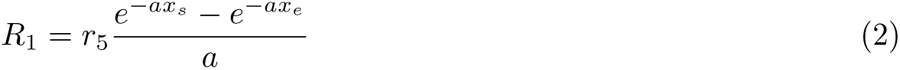

and

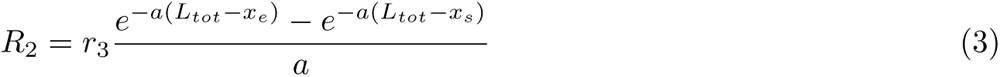

The model was fitted with non-linear least squares implemented in the ‘nls’ R function which returned predictions and standard errors. The goodness of fit was assessed by plotting observed vs. predicted values.

### Gradients of polymorphism

To carefully account for a correlation of recombination rates with levels of polymorphism we estimated both SNP density (SNP/kb) and 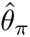 in each exon rank and gene classes in a similar manner as for recombination gradients. SNP density was the number of SNPs overlapping a given genomic feature found with ‘findOverlap’ from the GenomicRanges R package [82]. SNP density was averaged by calculating the mean per exon rank and gene classes. We also estimated 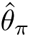 per site with ‘VCFtools’ on the final VCF dataset used for ‘LDhat’ [86]. The average 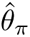 per exon rank and gene class was calculated as the sum of all 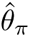 overlapping a given exon rank divided by the total number of sites covered by these exons.

## Acknowledgments

We wish to thank Laurent Duret, Pierre-Alexandre Gagnaire, Marie Raynaud, Julien Joseph, Nicolas Lartillot and all the other members of the HotRec ANR project for insighful discussions. Elise Rolland helped with the pipeline. We used the GENOUEST computing facility for bio-informatic analyses.

## Author contributions

Original idea: S.G.; Data analyses: T.B.; First draft: T.B.; Editing and revisions: T.B., S.G.

## Data and materials availability

Analyses scripts and documentation are available at https://github.com/ThomasBrazier/landrec-gradients. Data produced in this study are available at https://osf.io/3aekw/.

## Supplementary figures

**Figure S1:**
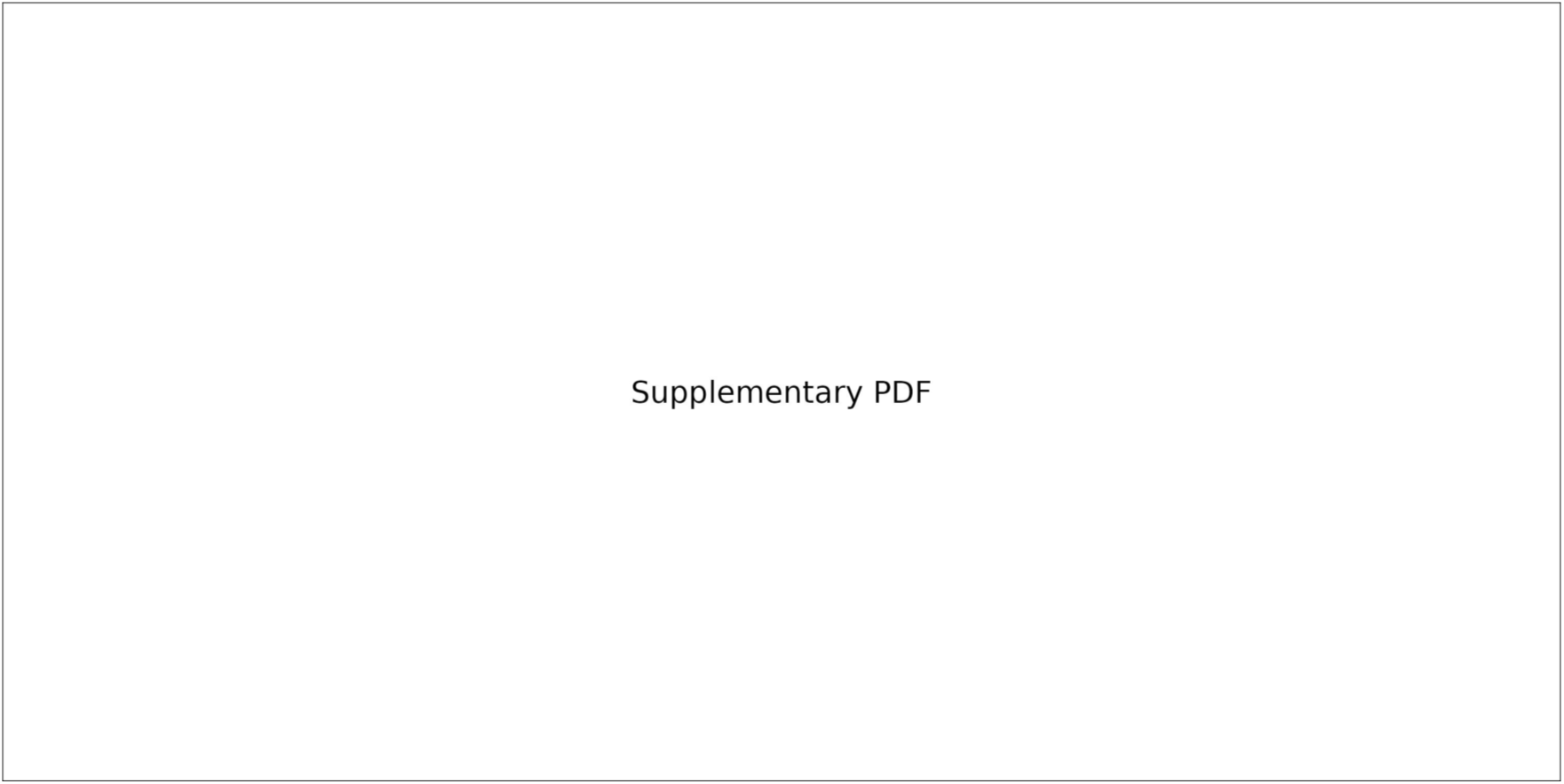
Chromosome recombination maps. (A) Fine-scale recombination maps estimated with LDhat. Mean *ρ*/kb for each inter-SNP interval. (B) Broad-scale recombination maps (cM/Mb, 100 kb windows) estimated from Marey maps in [1]. The grey ribbon is the 95% confidence interval estimated by 1,000 bootstraps on SNP markers.

**Figure S2:**
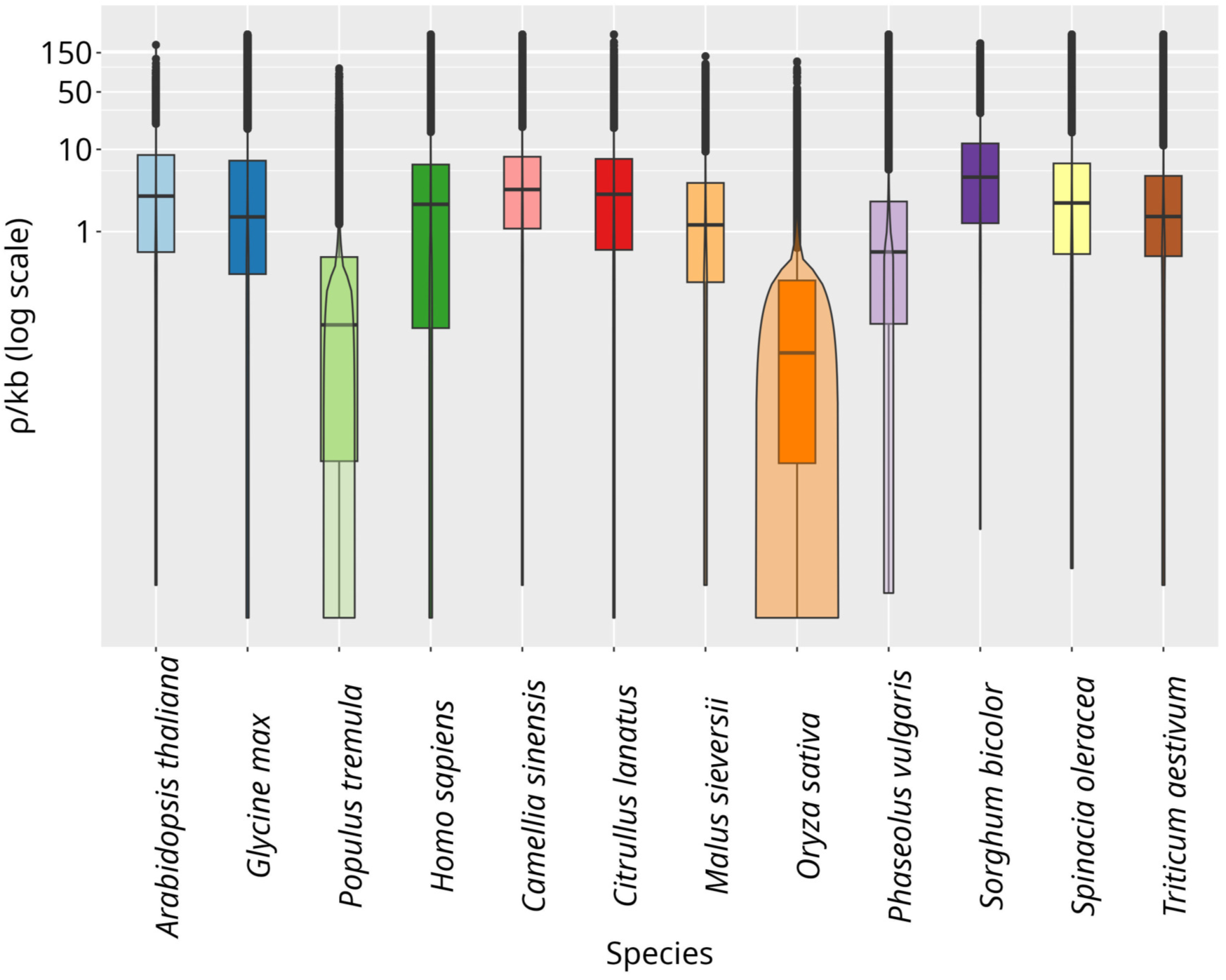
Species distributions of LDhat estimates (*ρ*/kb, log10 scale).

**Figure S3:**
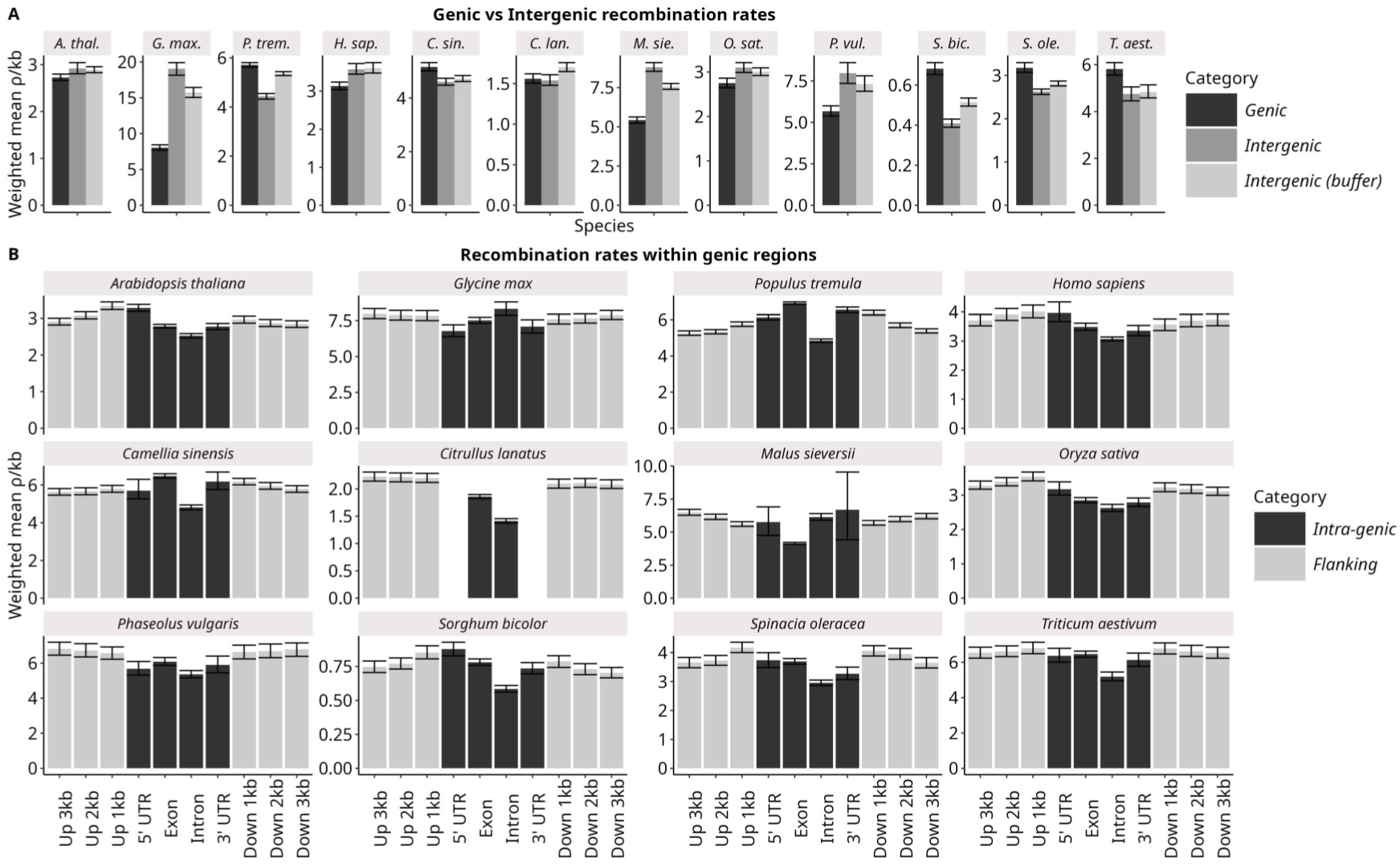
Variations in mean recombination rates (*ρ*/kb) around and within genes. (A) Recombination rates vary between genic and intergenic regions. In order to remove an effect of 5’ and 3’ regulatory regions at the proximity of genes, buffered intergenic regions were defined by excluding 3 kb flanking regions upstream and downstream of genes. (B) Within genic regions, the recombination rate varies between genomic features as well as in upstream and downstream flanking regions. The weighted mean (weighted by interval length) recombination rate and 95% confidence intervals were estimated by 1,000 bootstraps. Annotations of UTRs were not available for *Citrullus lanatus*.

**Figure S4:**
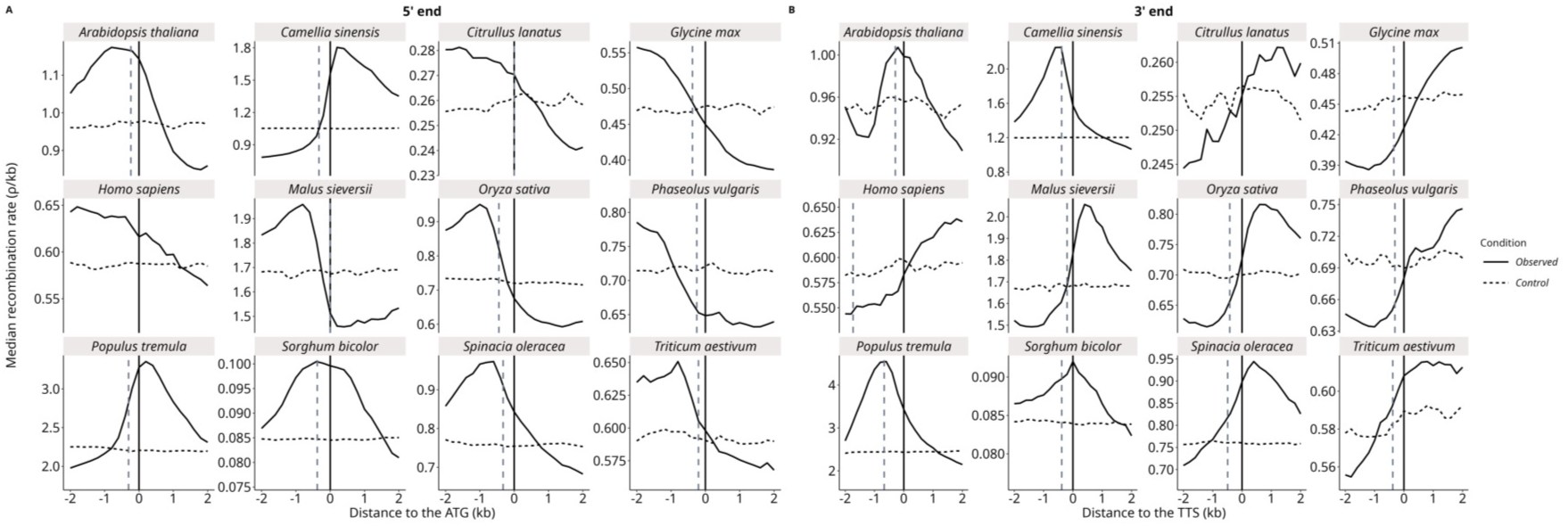
Recombination gradients along genes are mostly around Transcription Starting Sites (TSS) and Transcription Terminating Sites (TTS) among twelve species (including eleven plants). (A, B) The recombination rate (*ρ*/kb) was estimated in 200 bp windows as a function of the distance to the ATG start codon (start of first CDS) or the TTS and averaged among introns and exons. The vertical dashed black line is the mean distance to the TSS (i.e. the 5’ UTR) and the vertical dashed grey line is the mean 3’ UTR length. Random controls by randomly resampling intervals in the genome and re-scaled to the mean of the gradient.

**Figure S5:**
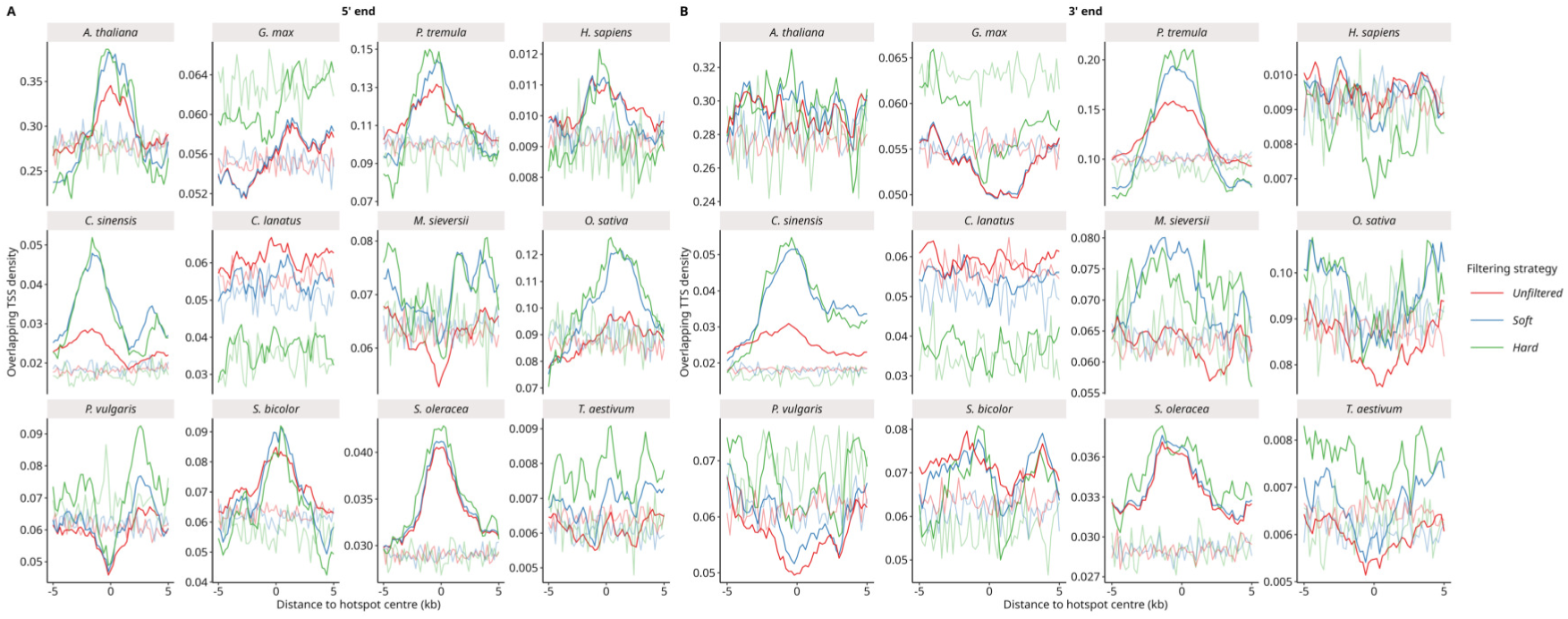
The density of TSS/TTS overlap around hotspot centre, expressed as a function of the distance to hotspot centre, for the three filtering strategies. For smoothing, TSS and TTS were defined as regions of 1 kb centered on the gene start and end position respectively. Soft lines are a control by randomly resampling hotspot intervals in a ± 50 kb neighboring region.

**Figure S6:**
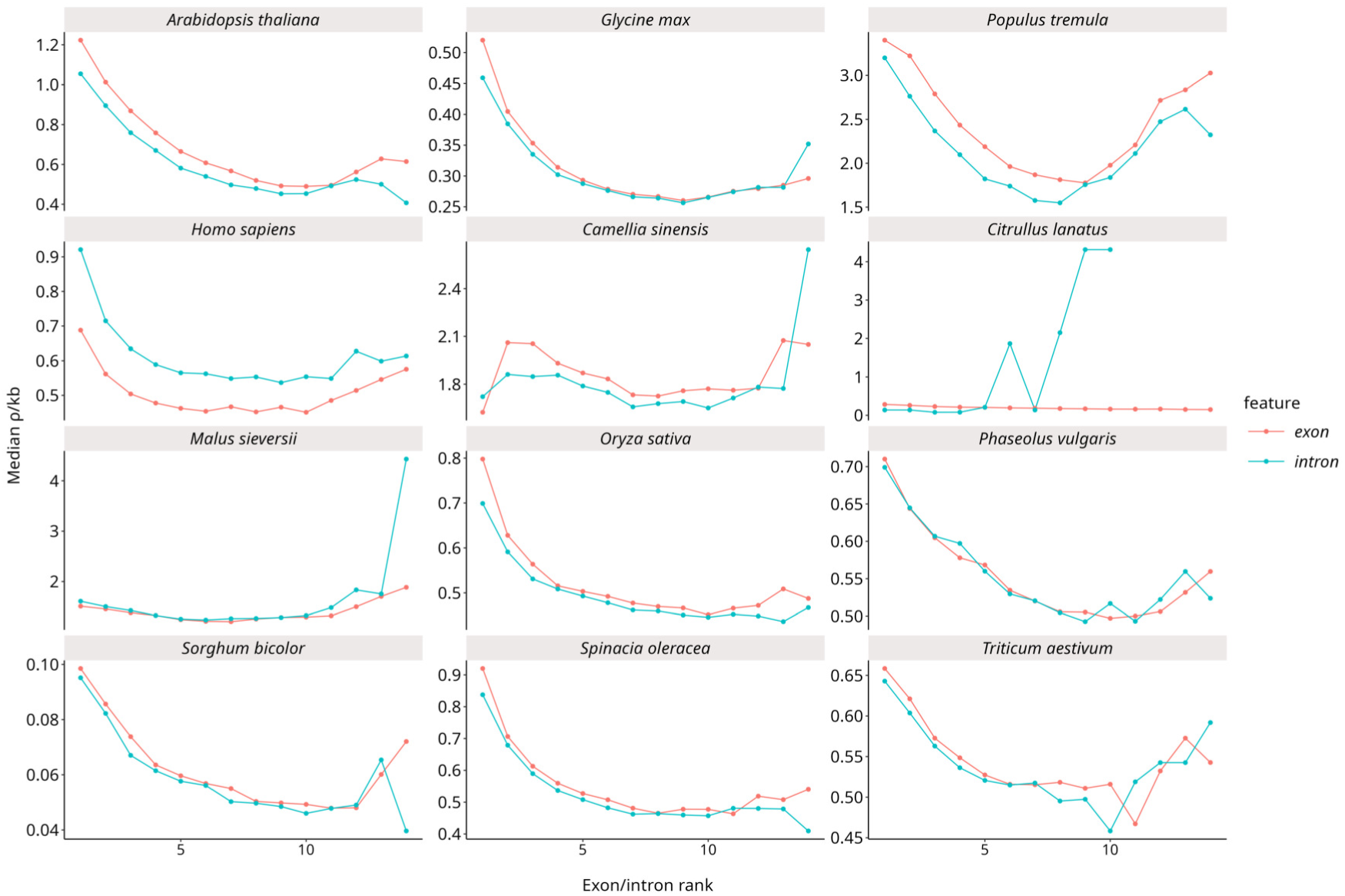
Genic recombination gradients of *ρ*/kb decomposed among introns and exons. The recombination rate (*ρ*/kb) was estimated in exons/introns as a function of exon/intron rank. Random controls by randomly resampling exon/introns intervals in the genome.

**Figure S7:**
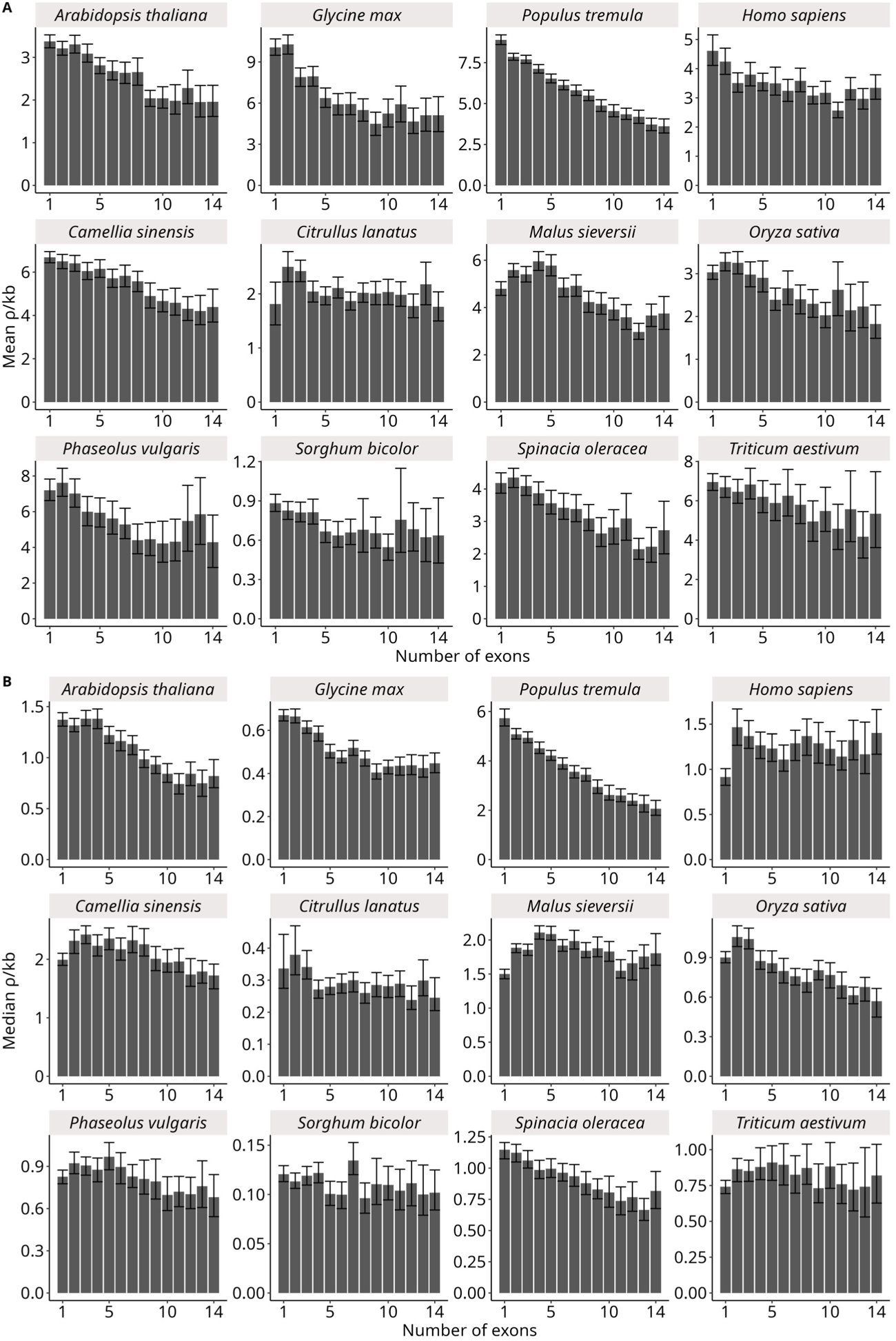
Weighted mean recombination rate (*ρ*/kb, weighted by the interval genomic size) as a function of gene size (i.e. number of exons). The 95% confidence intervals were estimated by 1,000 bootstraps.

**Figure S8:**
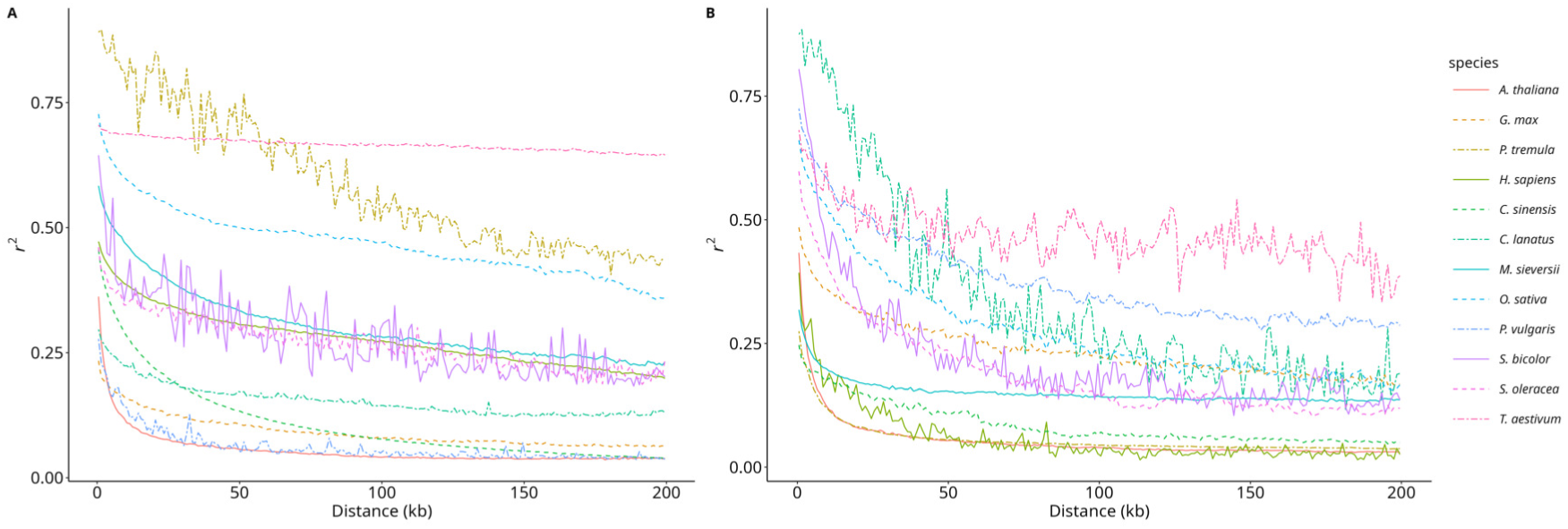
LD decay (*r*^2^) as a function of the pairwise distance between markers. (A) *r*^2^ measured genome wide. (B) *r*^2^ measured between SNPs in genic regions.

**Figure S9:**
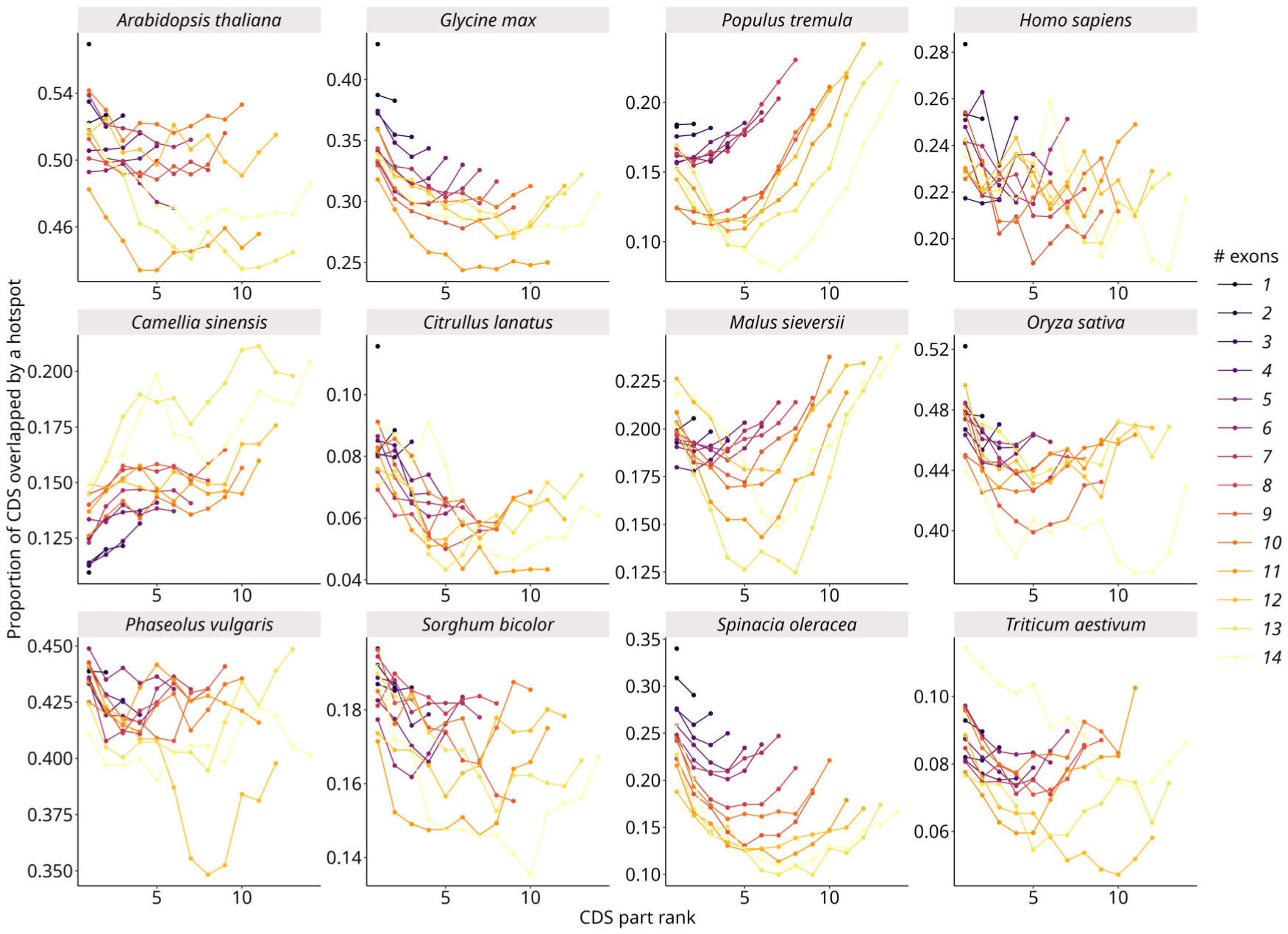
Gradients of hotspot overlap as a function of exon rank (CDS part). The proportion of CDS overlapping a hotspot in a given class of exon is the number of CDS overlapping at least one hotspot (soft filtered hotspots) divided by the total number of CDS of that class.

**Figure S10:**
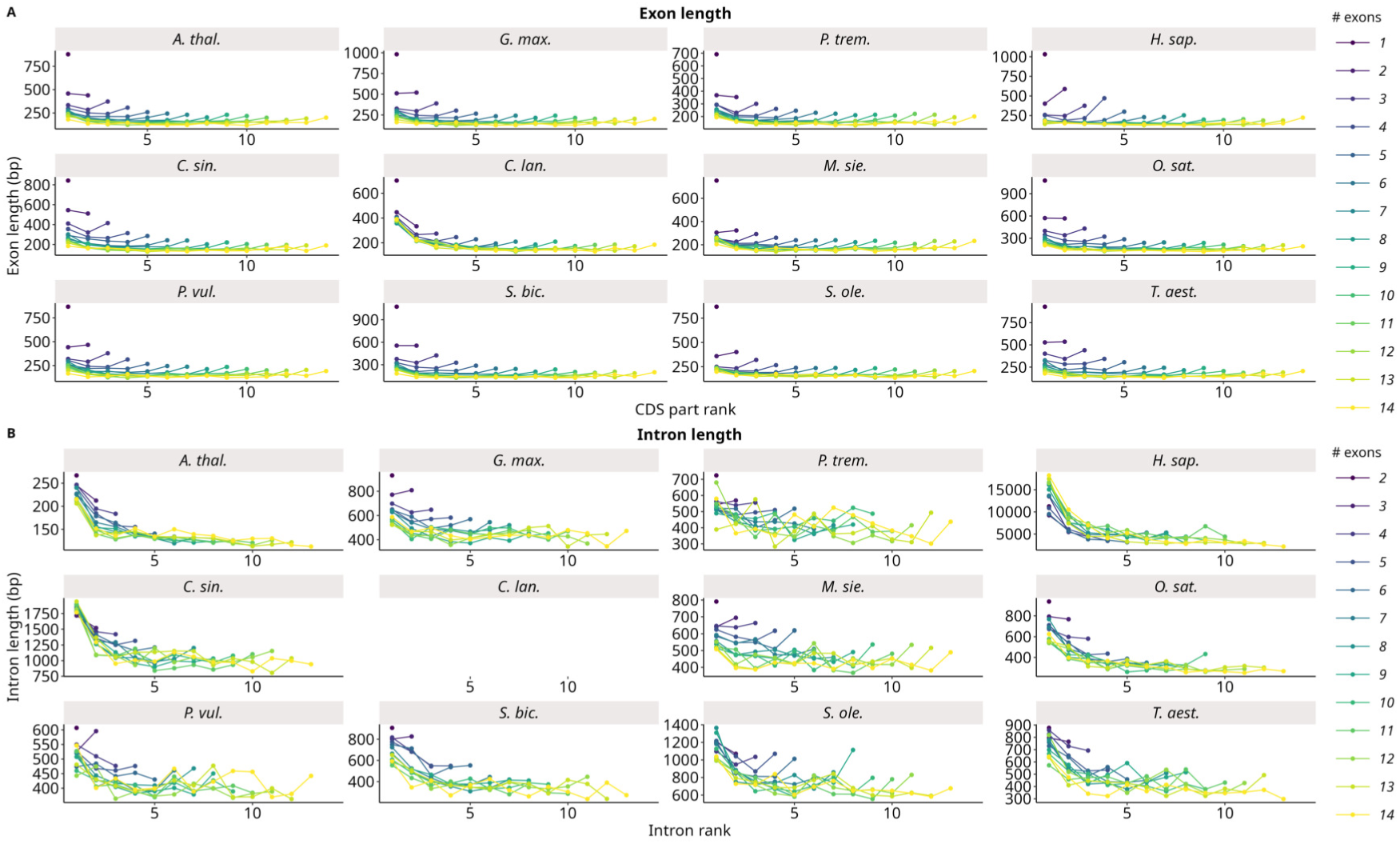
Exon and intron length along the coding sequence for twelve species. Exons/introns grouped per rank and gene size (number of exons). Only protein-coding genes were used. (A) Exon length. (B) Intron length. Introns in UTRs were not kept.

**Figure S11:**
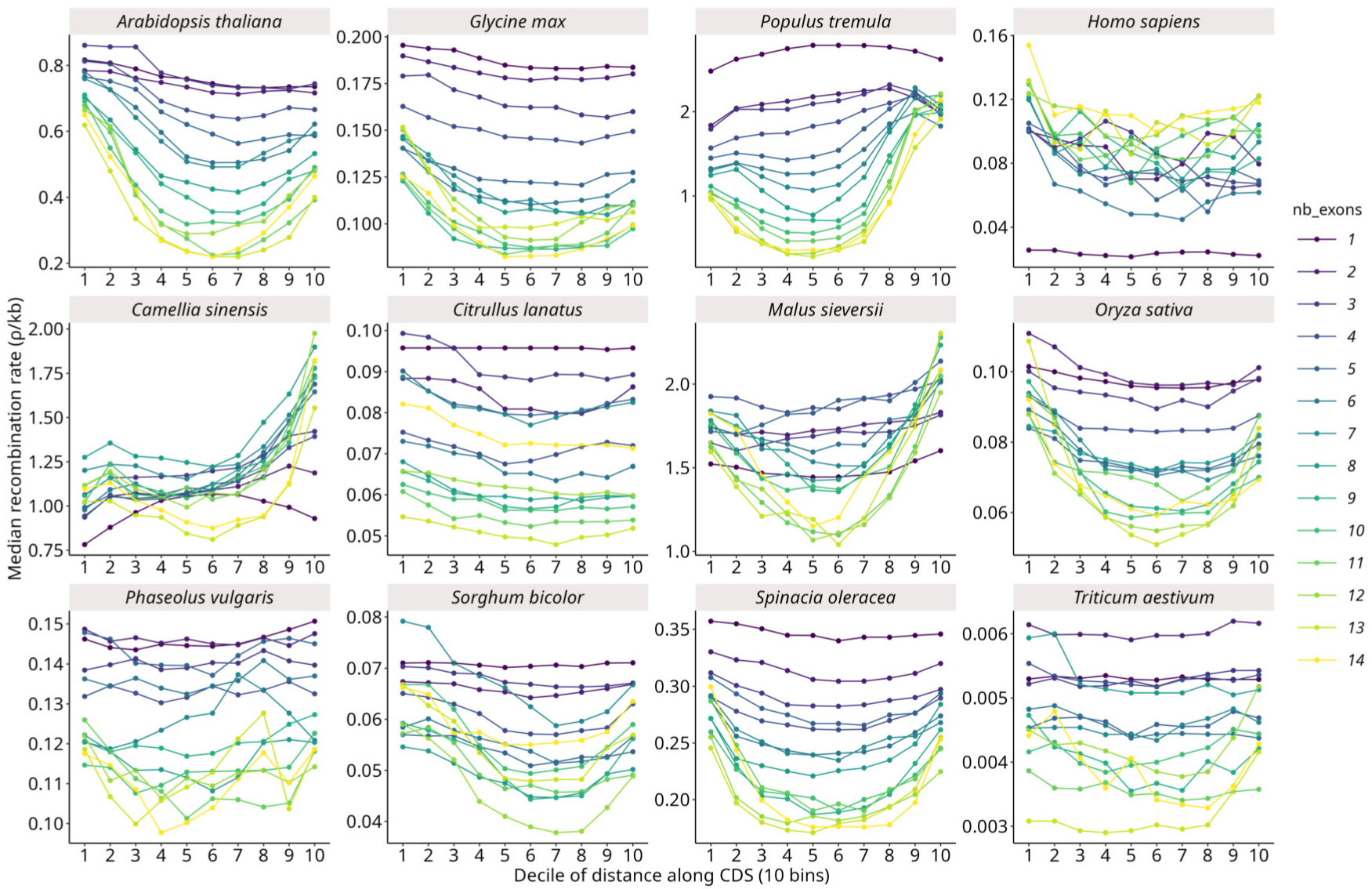
Recombination gradients per decile of distance across plant species. The recombination rate (median *ρ*/kb) was estimated in 10 deciles of distances as a function of their relative position along the gene (UTR + CDS) and genes were grouped by their number of exons. Only protein-coding genes were used.

**Figure S12:**
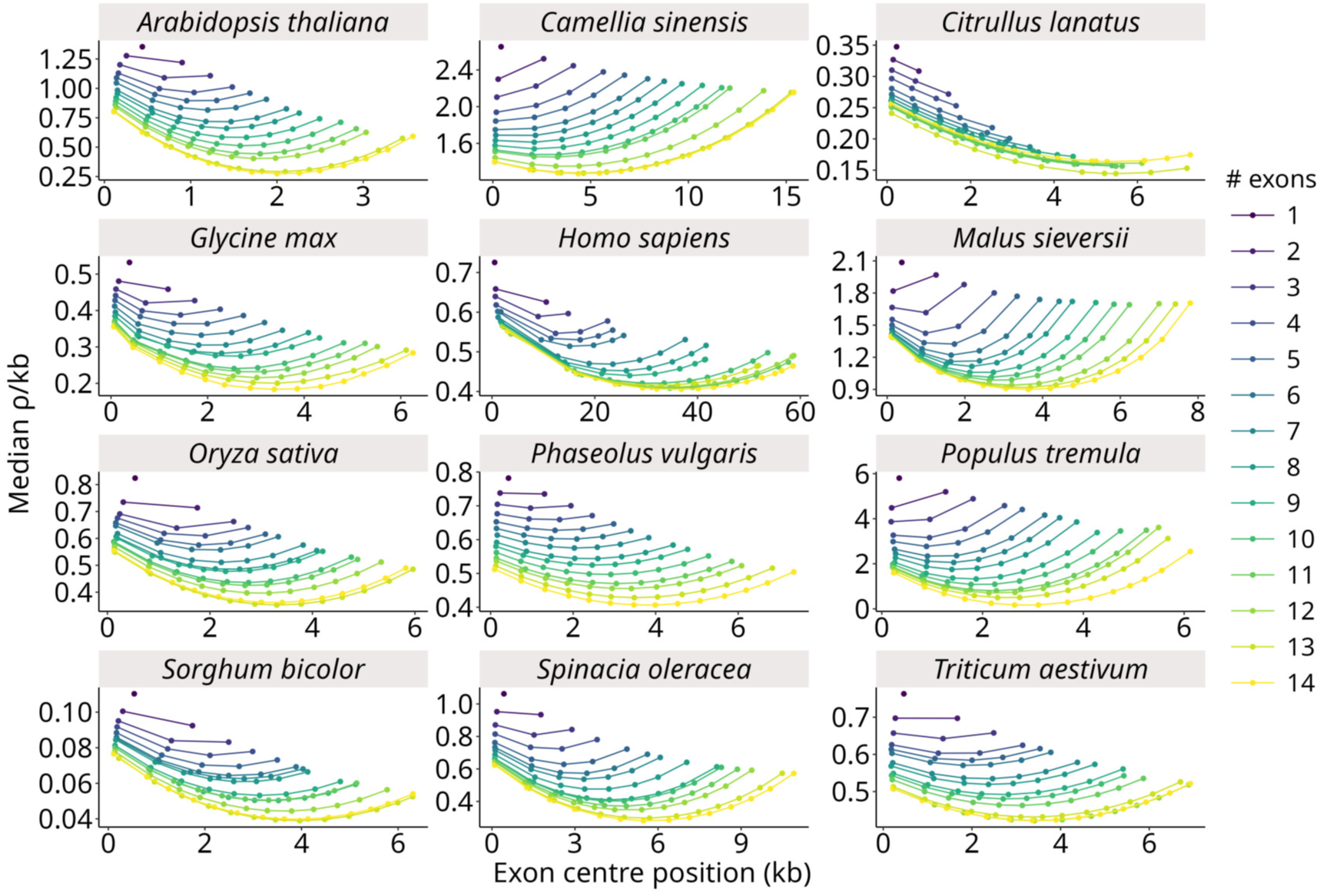
Gradients fitted with the hotspot model. Predicted *ρ*/kb as a function of exon genomic position (kb), pooled per exon rank and gene size (number of exons).

**Figure S13:**
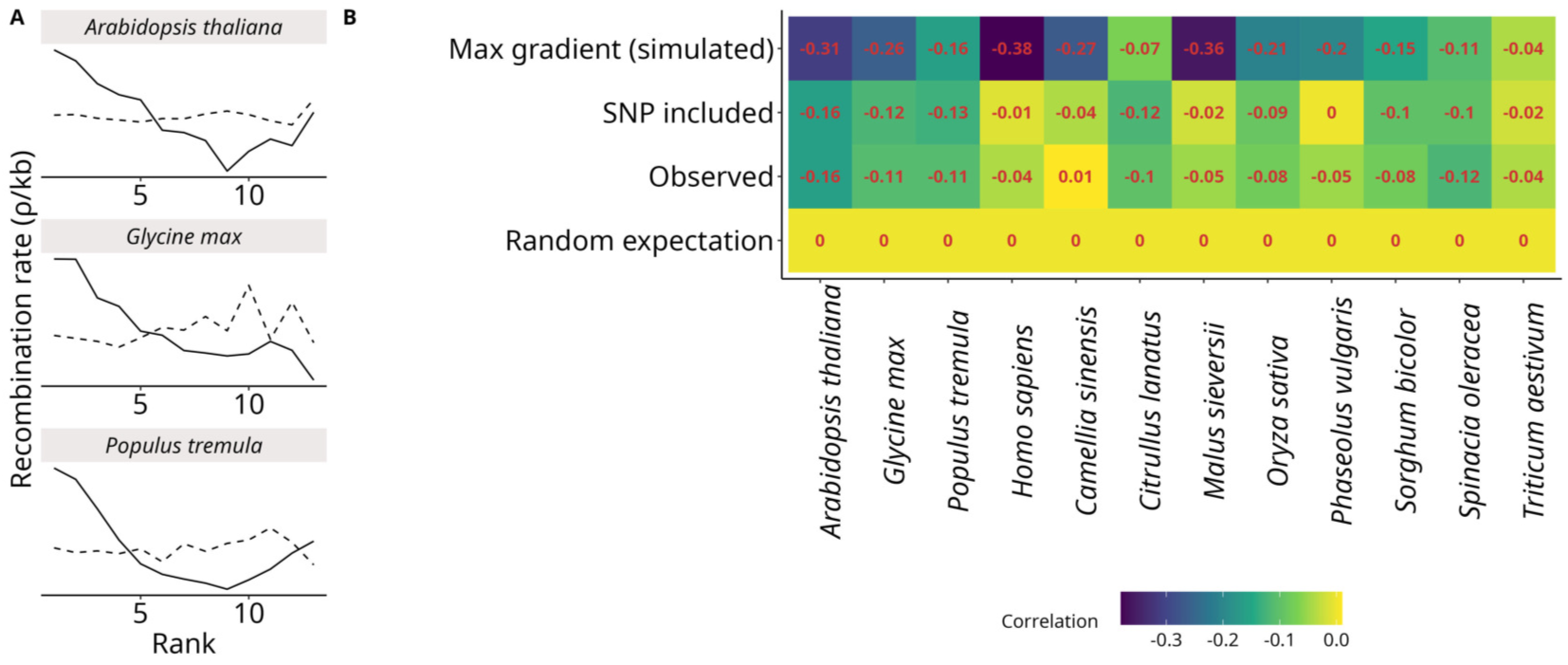
Evaluation of the performance of the method. (A) Strict gradients were estimated only with *ρ*/kb actually measured within the genomic interval. Random controls by resampling ranks (dashed lines). (B) Spearman’s correlation between *ρ*/kb and exon rank. From bottom to top, the random expectation was computed by resampling ranks with replacement; correlation in actual LD estimates (Observed); Strict gradient with only *ρ*/kb actually measured within the given exon (SNP included); a maximal ‘ideal’ gradient was simulated by reordering ranks by descending *ρ*/kb (Max gradient).

## Supplementary tables

Table S1: Metadata of the twelve datasets. Dataset name used during analyses, author of the original study, year of the original study, mating system, number of SNP before filtering, number of SNPs after filtering, total number of individuals in the dataset, number of individuals sampled for LDhat analyses, accession of the reference genome, number of chromosomes, number of genes, public database where to access original data (if relevant), direct link to the original data (if relevant), doi of the original study.

Table S2: Total number of genic vs intergenic hotspots for the three filtering strategies. Hotspots were considered as genic (intergenic, respectively) if they overlap (don’t overlap) a gene interval. The expected number of genic hotspots (± 95% confidence interval) was calculated by randomly resampling genic intervals.

Table S3: Sample size for each gene class (number of exons). Only genes with less than 15 exons were retained.

Table S4: Hyper-parameters used in the LDhat recombination pipeline, such as the sampled population, sampling, number of iterations and burnin size in the MCMC chain, pseudo-diploid procedure (0 for no, 1 for yes), the cutoff for runs of homozygosity in SMC++.

